# An atlas of fish genome evolution reveals delayed rediploidization following the teleost whole-genome duplication

**DOI:** 10.1101/2022.01.13.476171

**Authors:** Elise Parey, Alexandra Louis, Jérôme Montfort, Yann Guiguen, Hugues Roest Crollius, Camille Berthelot

## Abstract

Teleost fishes are ancient tetraploids stemming from an ancestral whole-genome duplication that may have contributed to the impressive diversification of this clade. Whole-genome duplications can occur via self-doubling (autopolyploidy) or via hybridization between different species (allopolyploidy). The mode of tetraploidization conditions evolutionary processes by which duplicated genomes return to a diploid state through meiosis resolution and subsequent genetic divergence (cytological and genetic rediploidization). How teleosts became tetraploid remains unresolved, leaving a fundamental gap to interpret their functional evolution. As legacy of the whole genome duplication, identifying orthologous and paralogous genomic regions across teleosts is challenging, hindering genome-wide investigations into their polyploid history. Here, we combine tailored gene phylogeny methodology together with a state-of-the-art ancestral karyotype reconstruction to establish the first high-resolution comparative atlas of paleopolyploid regions across 74 teleost genomes. We then leverage this atlas to investigate how rediploidization occurred in teleosts at the genome-wide level. We uncover that some duplicated regions maintained tetraploidy for over 60 million years, with three chromosome pairs diverging genetically only after the separation of major teleost families. This evidence suggests that the teleost ancestor was an autopolyploid. Further, we find evidence for biased gene retention along several duplicated chromosomes, contradicting current paradigms that asymmetrical evolution is specific to allopolyploids. Altogether, our results offer novel insights into genome evolutionary dynamics following ancient polyploidizations in vertebrates.

## Introduction

Since the first teleost fish genome sequence was published in 2002 (Aparicio et al. 2002), fish genomics has massively contributed to our understanding of vertebrate genome function and evolution. As an early-diverging vertebrate clade, teleosts are at an ideal phylogenetic position to conduct comparative analyses with tetrapods and study deep-rooting vertebrate features. Notably, the conservation of regulatory circuits and developmental pathways has turned zebrafish, medaka and - to a lesser extend - platyfish into model species for human diseases (Wittbrodt et al. 2002; Lieschke and Currie 2007; Schartl 2014). In addition, fish have become popular in evolutionary, ecological and physiological genomics, illuminating processes such as environmental adaptation, species diversification, social behavior or sex determination (Rittschof et al. 2014; Capel 2017; Salzburger 2018; Kim et al. 2019; Xie et al. 2019; Greenway et al. 2020). Around two hundred fish species have reference genome assemblies, and many more are expected to become available in the coming years (Rhie et al. 2021), requiring improved comparative frameworks to dissect the functional evolution of teleost genomes.

All teleost fish species are paleopolyploids: they descend from an ancient round of whole genome duplication (WGD), dated at approximately 320 Mya (Jaillon et al. 2004). This evolutionary event, referred to as the teleost-specific genome duplication (TGD), doubled all chromosomes and genes present in the teleost ancestor. The TGD has left a significant imprint on extant teleost genomes: while most duplicated genes have returned to a single-copy state, an important fraction of teleost genes remain in two copies, called ohnologs. For instance, 26% of all zebrafish genes are still retained as duplicated ohnologs (Howe et al. 2013). Evidence suggests that TGD duplicates have been involved in the evolution of innovations (Zakon et al. 2006; Moriyama et al. 2016; Escobar-Camacho et al. 2020), but it remains unclear how differential gene retention and functional divergence have sustained the impressive phenotypic diversity of the teleost clade.

WGD can arise through two mechanisms: autopolyploidization (within or between populations of a single species) or allopolyploidization (after hybridization of related species), with different consequences on subsequent genome evolution (Stebbins 1947; Mason and Wendel 2020). In particular, auto- and allopolyploidization differentially shape the rediploidization process, i.e. how the polyploid genome returns to a largely diploid state over time. Due to the initial sequence similarity between duplicated chromosomes (homeologs), young autopolyploid genomes are characterized by multivalent meiotic behaviour and tetrasomic inheritance, with recombination and gene conversion occurring between homeologous chromosomes. The mechanistic drivers of meiosis resolution remain poorly understood, with proposed roles for genomic rearrangements and reduced crossing-over frequency in promoting bivalent pairing (Bomblies et al. 2016; Mandáková and Lysak 2018). Maintenance of tetraploidy for millions of years over entire homeologs has previously been demonstrated in salmonids and Acipenseriformes (Robertson et al. 2017; Gundappa et al. 2021; Redmond et al. 2022). In these clades, rediploidization has been an extended process, with some duplicated genomic regions quickly restoring diploid behaviour, while other have maintained tetraploidy for tens of millions of years. In contrast, after allopolyploidization events, the extent of duplicated chromosome pairing depends on genetic similarity between the parental genomes. Localized homeologous exchanges, where sequences from one subgenome are substituted by sequences from the other, have been observed in allopolyploids, including allopolyploid crops and paleopolyploid carps (Lloyd et al. 2018; Li et al. 2021). These events however typically concern a minor fraction of allopolyploid genomes (less than 3%), and have never been shown to be maintained over millions of years.

The retention and divergence of gene copies is also affected by the nature of the polyploidization event. After allotetraploidization, one of the two subgenomes often loses more genes than the other (Garsmeur et al. 2014; Session et al. 2016; Cheng et al. 2018), although not always (Sun et al. 2017). This unequal gene retention has been linked to differences in transposable element repertoires and epigenetic silencing in the two subgenomes, and orients further functional evolution. Such differences can lead to reduced expression levels and relaxed selective pressure biased towards one of the subgenomes, which then accumulates more gene losses (Freeling et al. 2012; Bird et al. 2021). Conversely, in the case of autopolyploidy, gene losses are expected to affect both homeologs equally due to their high similarity. It remains unclear whether the teleost whole-genome duplication corresponds to an ancestral auto- or an allotetraploidization event, an important gap in our understanding of the early vertebrate evolutionary processes that led to the diversification of teleosts (Martin and Holland 2014; Conant 2020).

Importantly, the redundancy in fish genomes can be appreciated at the macrosyntenic level, where remnants of ancestrally duplicated chromosomes form runs of large duplicated regions known as double-conserved syntenic regions (DCS) (Postlethwait et al. 2000; Taylor et al. 2003; Jaillon et al. 2004). The precise identification and delimitation of teleost DCS regions is the key to reconstructing how rediploidization occurred and its associated impact on teleost evolution. However, WGDs present severe challenges to both gene phylogeny and ancestral genome reconstruction methodologies (Nakatani and McLysaght 2017; Zwaenepoel and Van de Peer 2019; Parey et al. 2020). Previous characterizations of DCS in teleosts were therefore limited to regions of highly conserved gene order or small species sets: the largest multi-species dataset comprised eight fish species and included ∼29% of all genes in DCS (Conant 2020).

Here, we apply a phylogenetic pipeline specifically developed for WGD events (Parey et al. 2020) to retrace the evolutionary history of the genes and chromosomes of 74 teleost species encompassing most of the major fish clades. We combine these phylogenetic trees with the latest ancestral reconstruction of the pre-TGD ancestral karyotype (Nakatani and McLysaght 2017) to build a comprehensive comparative atlas of TGD-duplicated regions in teleost fish genomes. We then leverage this comparative atlas of teleost genomes to reconstruct how rediploidization occurred genome-wide following the teleost genome duplication, revealing how the ancestral teleost became tetraploid.

## Results

### Construction of TGD-aware teleost gene trees

We collected a dataset of 101 vertebrate genomes, including 74 teleost fish, 2 non-teleost fish (bowfin and spotted gar, which did not undergo the teleost genome duplication or TGD), 20 other vertebrates of which 6 are mammals, and 5 non-vertebrate genomes (Supplementary Figure S1; Supplementary Table S1). We used TreeBeST to reconstruct the phylogenetic relationships of 26,692 gene families across those 101 genomes (Methods; (Vilella et al. 2009; Herrero et al. 2016)). We then applied SCORPiOs to correctly place the TGD in these gene trees, using the bowfin and the spotted gar as reference outgroups (Parey et al. 2020). Briefly, SCORPiOs leverages synteny information to distinguish WGD-descended orthologs from paralogs when sequence information is inconclusive. After a WGD, orthologous genes are expected to be embedded in orthologous neighborhoods. For each individual gene tree, SCORPiOs “crowd-sources” additional information from local syntenic genes to identify errors in orthology relationships. SCORPiOs then reorganizes those gene trees to accurately position the WGD duplication node, if the synteny-consistent solution is equally supported by the sequence alignment. These 26,692 WGD-aware teleost gene trees predict that the ancestral genome of teleost fish contained 46,206 genes after the duplication event, in line with the latest estimates from the Ensembl database (49,255 ancestral teleost genes in release v100; Methods).

### A high-resolution atlas of the TGD duplication across 74 teleost genomes

Teleost fish genomes are mosaics of duplicated regions, formed through rearrangements of duplicated ancestral chromosomes (Supplementary Figure S2A). Long-standing efforts have been made to reconstruct the ancestral teleost karyotype before the whole-genome duplication (Jaillon et al. 2004; Kasahara et al. 2007; Nakatani and McLysaght 2017). According to the state-of-the-art reference, this ancestral teleost karyotype comprised 13 chromosomes (Nakatani and McLysaght 2017). Nakatani and McLysaght delineated between 353 and 690 megabase-scale genomic regions that descend from these 13 ancestral chromosomes in zebrafish, tetraodon, stickleback and medaka.

We combined this pre-TGD ancestral karyotype with our gene trees to identify regions and genes that descend from sister duplicated chromosomes across all 74 teleosts in our dataset (Methods; Supplementary Figure S2). First, we transformed the reference segmentations of the zebrafish, tetraodon, stickleback and medaka genomes (reference species) from 13 to 26 ancestral chromosomes after the duplication (1a, 1b, …, 13a, 13b). In each genome, we iteratively grouped regions from an ancestral chromosome into two post-duplication copies by minimizing intragroup paralogs (Methods; Supplementary Figure S2B). To assess robustness, we performed 100 groupings with random restart and found that genes were assigned to the same chromosome copy in 80% of iterations on average. We then identified orthologous ancestral chromosomes across all four species using gene orthologies, and arbitrarily named one of each pair ‘a’ and ‘b’ (Supplementary Figure S2C). Next, we propagated these ancestral chromosome annotations to the other 70 teleost genomes using gene orthology relationships (Supplementary Figure S2D). For 1,303 gene trees (7%), orthologs from the four reference species were not assigned to the same ancestral chromosome, and we resolved these inconsistencies by assigning all orthologous genes, including those of the reference species, to the most represented ancestral chromosome. This process results in a 74-species comparative genomic atlas with genes annotated to post-duplication chromosomes, along with fully resolved orthology and paralogy links between all included species (Supplementary Figure S2E).

This comparative atlas integrates 70% to 90% of each genome into 24,938 post-duplication gene families (Figure 1A-B), and greatly improves upon the state-of-the-art both in terms of species and genomic coverage. The atlas reveals that teleost fishes vary substantially in their retention of duplicated gene copies (ohnologs) since the TGD, which make up 33% of the arowana genome but only 19% of the cod genome (Supplementary Table S2). In general, Osteoglossiformes, Otomorpha and Salmoniformes species have retained more ohnologs from the TGD than other Euteleosteomorpha clades that diverged later. The atlas is available on the Genomicus database webserver (see Data availability and implementation).

**Figure 1:**
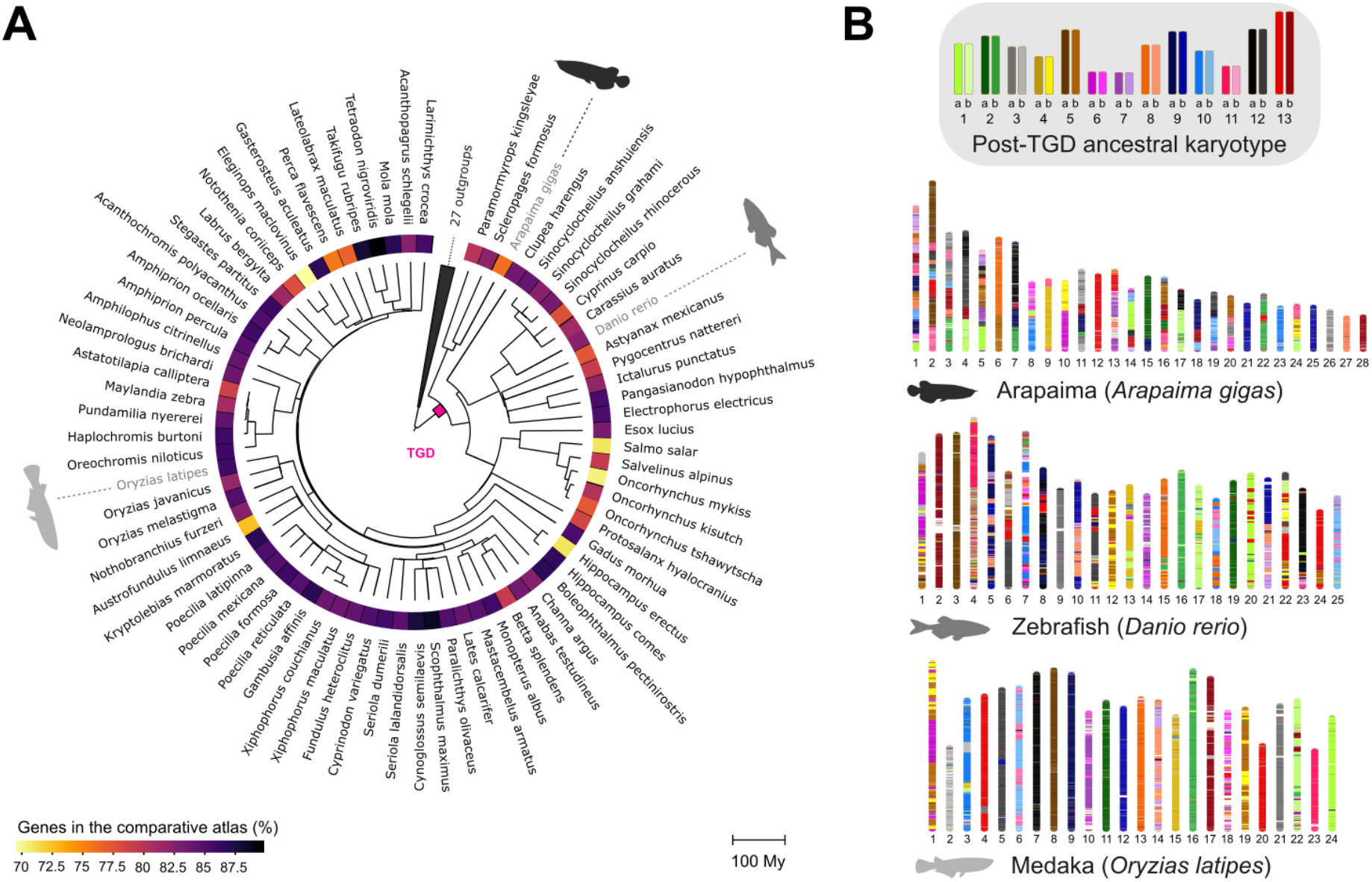
Comparative atlas of WGD-duplicated regions across 74 teleosts. **A**. Phylogenetic tree of the 74 teleost genomes in the comparative atlas and 27 outgroups. The color map represents the proportion of genes from each species annotated in the comparative atlas. Divergence times were extracted from TimeTree (Kumar et al. 2017). **B**. Karyotype paintings using the comparative atlas. At the top, we show the inferred ancestral karyotype after the teleost whole-genome duplication (TGD). Below, karyotypes of three teleost genomes are colored by their ancestral chromosome of origin according to the comparative atlas (1a, 1b, …, 13a, 13b).

### Quality checking the teleost genome comparative atlas

As a quality check, we assessed discrepancies between the pre-TGD karyotype reconstruction by (Nakatani and McLysaght 2017) and our ancestral chromosome assignations. Both should be globally congruent because the ancestral karyotype provides the groundwork for the comparative atlas, but discrepancies can arise when orthologous genes are assigned to different pre-duplication chromosomes between the four reference species. In this case, we resolve the inconsistency by reassigning all orthologs to the most represented pre-duplication chromosome, which will be different from the original assignation in (Nakatani and McLysaght 2017) for at least one species. We found that 9% of zebrafish genes differ in ancestral chromosome assignation between our comparative map and the pre-TGD karyotype, versus 2-3% in medaka, tetraodon or stickleback. This likely reflects small-scale rearrangements in zebrafish captured by our individual gene trees but missed by the macrosyntenic approach of (Nakatani and McLysaght 2017). Alternatively, zebrafish genes may be more frequently placed incorrectly in our gene trees and assigned to the wrong orthology group, which may lead to erroneous ancestral chromosome reassignations. To explore this possibility, we identified gene trees that remain synteny-inconsistent after correction by SCORPiOs (i.e. whose orthology relationships are inconsistent with those of their surrounding genes), and therefore potentially contain orthology errors. Zebrafish genes reassigned to a different ancestral chromosome in our comparative map are not over-represented in synteny-inconsistent trees (7% vs 14% for all zebrafish genes), suggesting that their orthology relationships and chromosomal reassignations are overall well-supported by their sequences and syntenic gene neighborhoods.

Additionally, we used random noise simulations to demonstrate that the comparative atlas is robust to potential uncertainty or errors in the original ancestral genome reconstruction. The delineation of genomic regions descended from each pre-duplication chromosome from (Nakatani and McLysaght 2017) relies on the identification of conserved synteny blocks and their breakpoints between teleost genomes and outgroups. Because breakpoint locations are challenging to determine - as also attested by lower posterior probabilities close to breakpoints in (Nakatani and McLysaght 2017) - the region boundaries can vary in precision. We shifted the positions of these boundaries with increasingly large errors, mimicking situations where up to 25% of genes change pre-duplication chromosome assignations in each of the reference genomes (Methods). We found that even large errors in region boundaries did not majorly affect the final atlas, with only 11% of genes changing ancestral chromosome assignations at the highest noise settings (Supplementary Figure S3).

Finally, we report that correcting gene trees with SCORPiOs had a decisive impact on the establishment of the comparative atlas, enabling the inclusion of a significantly larger fraction of teleost genes (83% vs 61%, Supplementary Figure S4). The teleost comparative genomic atlas represents therefore a reliable, comprehensive and robust resource for fish genomics.

### The teleost duplication was followed by delayed rediploidization

The comparative genomic atlas is the first resource that allows an in-depth, genome-wide analysis of the TGD, and more importantly of the early genome evolution processes that followed after the TGD. Previous work has revealed that auto- and allotetraploids significantly differ in their early evolution (Stebbins 1947; Garsmeur et al. 2014; Mason and Wendel 2020): specifically, autotetraploids initially harbour four near-identical chromosome sets, and therefore can maintain meiotic recombination and tetrasomic inheritance for tens of million years. Previous work in salmonids and Acipenseriformes have revealed that meiosis resolution is dynamic and occurs in waves across homeologous regions (Robertson et al. 2017; Gundappa et al. 2021; Redmond et al. 2022). Sequence exchanges due to prolonged recombination are detectable because they delay the divergence of duplicated regions until after meiosis is fully resolved, resulting in different phylogenetic expectations regarding ohnolog sequences evolution. Depending on the respective timings of rediploidization and speciation, ohnologs can either follow the AORe (Ancestral Ohnolog Resolution) or LORe (Lineage-specific Ohnolog Resolution) models (Figure 2A), introduced in (Robertson et al. 2017). In the AORe model, meiosis is resolved before lineages split, thus initiating ohnologous sequence divergence before speciation. In the LORe model, because recombination still persists when speciation occurs, ohnologs share more sequence similarity within clades than across clades, and can therefore be mis-identified as clade-specific duplications.

**Figure 2:**
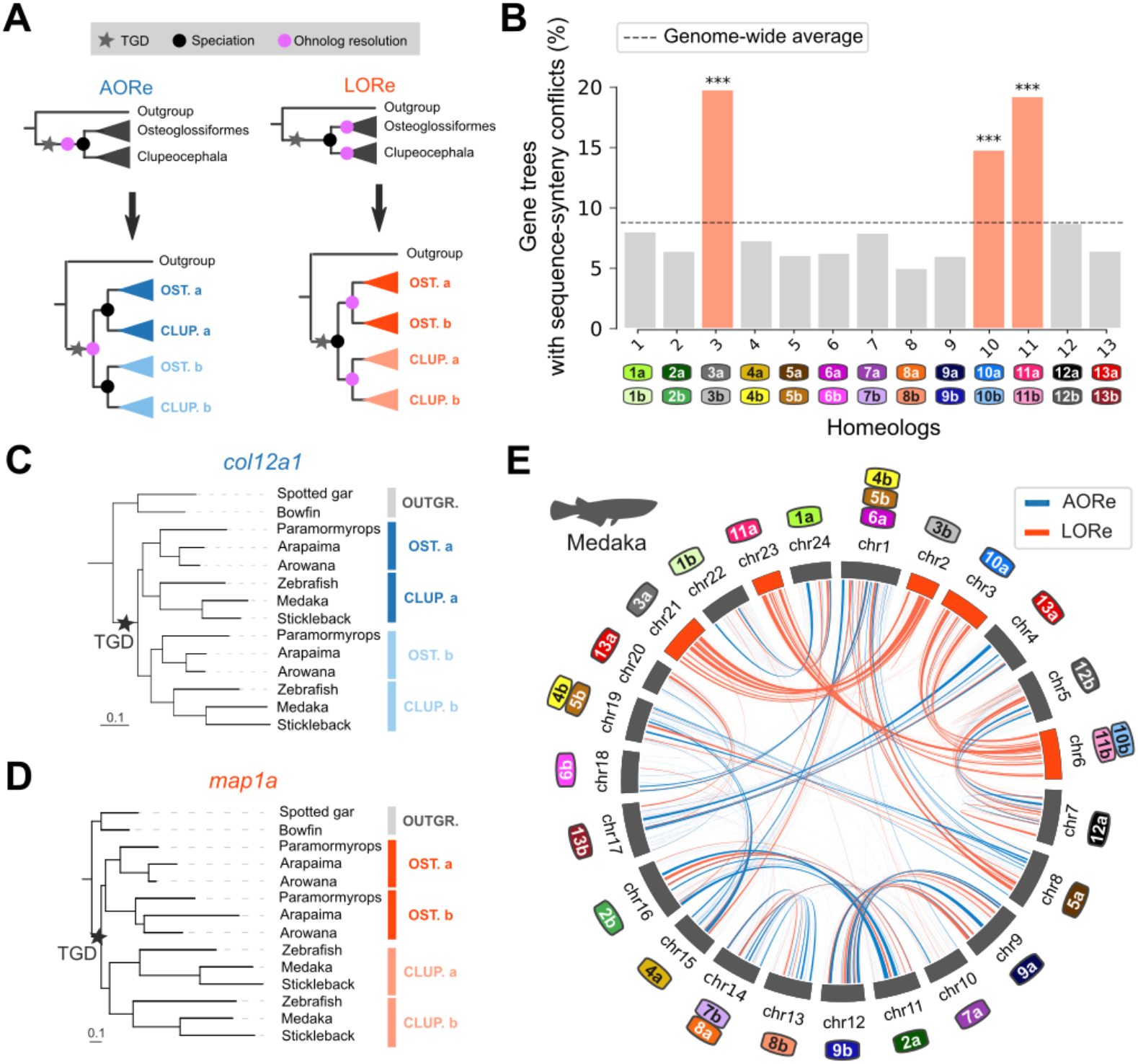
Delayed rediploidization following the TGD. **A**. Gene tree topologies expected under the AORe and LORe models. The AORe tree topology assumes that rediploidization was complete before the divergence of Osteoglossiformes and Clupeocephala, initiating ‘a’ and ‘b’ gene sequence divergence before speciation. The LORe tree topology assumes that rediploidization was completed only after the divergence of Osteoglossiformes and Clupeocephala, delaying ‘a’ and ‘b’ duplicated sequences divergence to after speciation. **B**. Ancestral chromosomes 3, 10 and 11 are enriched in sequence-synteny conflicts (Methods, *** p < 0.001, hypergeometric tests with Benjamini-Hochberg correction for multiple testing). Color labels identify ancestrally duplicated chromosomes as in Figure 1A. **C**. Examples of an AORe gene tree. For the *col12a1 - a-col12a1b* family, the LORe topology is inconsistent with gene sequence evolution (p = 4e-09, AU-test). **D**. Example of a LORe gene tree. For the *map1aa - map1ab* family, AORe was rejected (p-value = 0.001, AU-test). **E**. AORe and LORe gene families visualized on the medaka karyotype. Medaka chromosomes are annotated as numbers, while color labels represent ancestral chromosomes (Methods), as in (B). Homeologs 3, 10 and 11 almost entirely rediploidized later than the Osteoglossiformes/Clupeocephala divergence.

To investigate the polyploidization mode and rediploidization processes of teleost fishes, we established whether meiosis was fully resolved with bivalent chromosomal pairing in the teleost ancestor before extent teleost lineages diverged, or whether some duplicated genomic regions still recombined during meiosis at that time. We focus our analysis at the Osteoglossiformes and Clupeocephala lineage split, which is the earliest possible speciation point in the teleost phylogeny (Parey et al. 2022), dated approximately at 267 Mya (Kumar et al. 2017). We selected a subset of six teleost genomes *(Paramormyrops*, arapaima, Asian arowana, zebrafish, medaka and stickleback) to ensure an equal representation of Osteoglossiformes and Clupeocephala genomes in the dataset, along with outgroup genomes (Methods, Supplementary Figure S5). We developed an extension to SCORPiOs named LORelEi for “<u>L</u>ineage-specific <u>O</u>hnolog <u>Re</u>solution <u>E</u>xtension”. SCORPiOs LORelEi is built around two modules (“diagnostic” and “likelihood tests”) to identify delayed rediploidization in gene trees. The LORelEi diagnostic module leverages the gene tree correction applied by SCORPiOs to identify sequence-synteny conflicts. Sequence-synteny conflicts arise when the orthology relationships of a gene family are inconsistent with conserved synteny information, but correcting the WGD duplication node to rectify this inconsistency induces a significant drop in likelihood estimated from sequence divergence. We grouped gene trees with sequence-synteny conflicts according to their ancestral chromosome of origin in the comparative atlas. Three anciently duplicated chromosome pairs (3, 10 and 11) are significantly enriched in sequence-synteny conflicts, thus hinting towards prolonged recombination between these homeologs after the TGD (Figure 2B; p < 0.05, hypergeometric tests corrected for multiple testing).

We next sought to confirm that the identified sequence-synteny conflicts were consistent with the phylogenetic expectations of delayed rediploidization. We used the LORelEi likelihood test module to explicitly compare the likelihoods for the tree topologies expected under the AORe and LORe rediploidization models, where ohnolog resolution occurs before or after the Osteoglossiformes/Clupeocephala lineage split, respectively (Figure 2A; Methods). We performed these likelihood tests on 5,557 gene trees which retain both ohnologs in at least one of the descending lineages. For 638 gene trees, the early resolution topology (AORe) was significantly more likely, while the late resolution topology (LORe) was favored for 1,361 trees (likelihood AU-tests, alternative topology rejected at α=0.05; no significant differences for the remaining 3,558 trees). For example, the *col12a1a - col12a1b* ohnologs had stopped recombining and accumulated independent substitutions by the time Osteoglossiformes and Clupeocephala diverged (Figure 2C). On the other hand, the *map1aa - map1ab* TGD ohnologs presumably still recombined when the taxa diverged, and both ohnologs diverged independently later on in each lineage (Figure 2D). We mapped the location of these genes on the medaka karyotype, which remains close to the ancestral teleost karyotype, to identify chromosomal regions of ancestral and lineage-specific rediploidization (Figure 2E; Methods). This visualization revealed that ancestral chromosome pairs 3, 10 and 11 had not yet initiated sequence divergence and were likely still recombining as tetrads when teleosts started diversifying, with large runs of LORe ohnologous families spanning the entire length of their descendant chromosomes in medaka (chromosomes 2, 3, 6, 21 and 23). The other homeologs appear as a mix of localized AORe and LORe regions, suggesting that rediploidization was ongoing.

This snapshot of the rediploidization status at the time of the Osteoglossiformes/Clupeocephala lineage split shows that rediploidization was not uniform across the ancestral teleost genome, consistent with results in salmonids (Robertson et al. 2017; Gundappa et al. 2021). In salmonids, different waves of rediploidization have been linked with variations in gene functions (Gundappa et al. 2021). In teleosts, we found no evidence of functional enrichments in either the AORe genes, which rediploidized early, or the LORe genes carried by late-rediploidized ancestral chromosomes 3, 10 and 11. Interspersed LORe genes on other homeologs were functionally enriched for different pathways, including “TGF-beta signaling pathway”, “regulation of transferase activity” and “regulation of intracellular signal transduction” (corrected p-values< 0.05; Supplementary Table S3, S4; Methods). One possible explanation to the functional differences between the two sets of LORe genes might be that homeologous chromosomes 3, 10 and 11 remained tetraploid due to specificities in their nuclear organization and topological features which prevented rediploidization; while sequence exchanges at LORe loci on other homeologs may have been locally maintained due to selection, for example by removing deleterious deletions mutations or transferring favourable alleles at all gene loci. In this speculative model, meiosis resolution would largely occur in waves because some chromosomes or chromosomal regions are mechanistically “easier” to rediploidize than others.

Altogether, we provide the first evidence that entire chromosomes experienced delayed rediploidization in teleosts and continued to exchange genetic material between homeologs for at least 60 million years after the teleost whole genome duplication. This prolonged exchange of genetic material between duplicated chromosomes after the TGD strongly suggests that the teleost ancestor was an autotetraploid.

### The teleost genome experienced biased gene retention after duplication

Previous observations on a limited subset of genes have suggested that the ancestral teleost genome has experienced biased gene retention after the TGD (Conant 2020). Biased gene retention is generally considered a hallmark of allopolyploidization (Garsmeur et al. 2014), but has mostly been observed in plants, and the mechanisms of gene retention in polyploid vertebrate genomes may differ. We tested whether the ancestral tetraploid teleost underwent biased gene retention, with one duplicated chromosome copy systematically retaining more genes than the other. We computed gene retention in non-overlapping 10-gene windows along duplicated chromosome copies in several teleost genomes, using an outgroup fish as proxy for the ancestral gene order (Figure 3A-B, Supplementary Figures S6-S7, Methods) and total gene retention on homeologs in the ancestor (*Osteoglossocephalai*) of all 74 teleosts (Supplementary Table S5). We uncover a consistent, significant bias in gene retention on ancestral chromosome pairs 3, 4, 7, 11 and 13, where genes were preferentially retained on one homeolog over the other, regardless of study species. We find additional but weaker evidence for biased gene retention on ancestral chromosome pairs 2, 5, 6, 8 and 9 in some combinations of genome comparisons. We however do not find evidence of retention bias between ancestrally duplicated copies for chromosome pairs 1, 10, 12. This preferential gene retention is not an artifact of the atlas construction due to genes unassigned to either duplicated chromosome: indeed, conservatively assigning such genes to the homeolog with the lowest retention did not offset the retention imbalance (Supplementary Table S6). In summary, we find that genes were preferentially retained on one duplicated chromosome in a subset of ancestral chromosomes, a character more frequently observed in allopolyploids (Garsmeur et al. 2014; Cheng et al. 2018). Of note, the chromosome copy with highest gene retention is typically annotated as ‘a’ in the atlas as a consequence of the construction process. As the TGD is likely an autopolyploidization event, ‘a’ and ‘b’ correspond to different chromosome copies and are interchangeable for each chromosome pair – ‘a’ and ‘b’ chromosomes should not be interpreted as distinct parental subgenomes of allopolyploidization.

**Figure 3:**
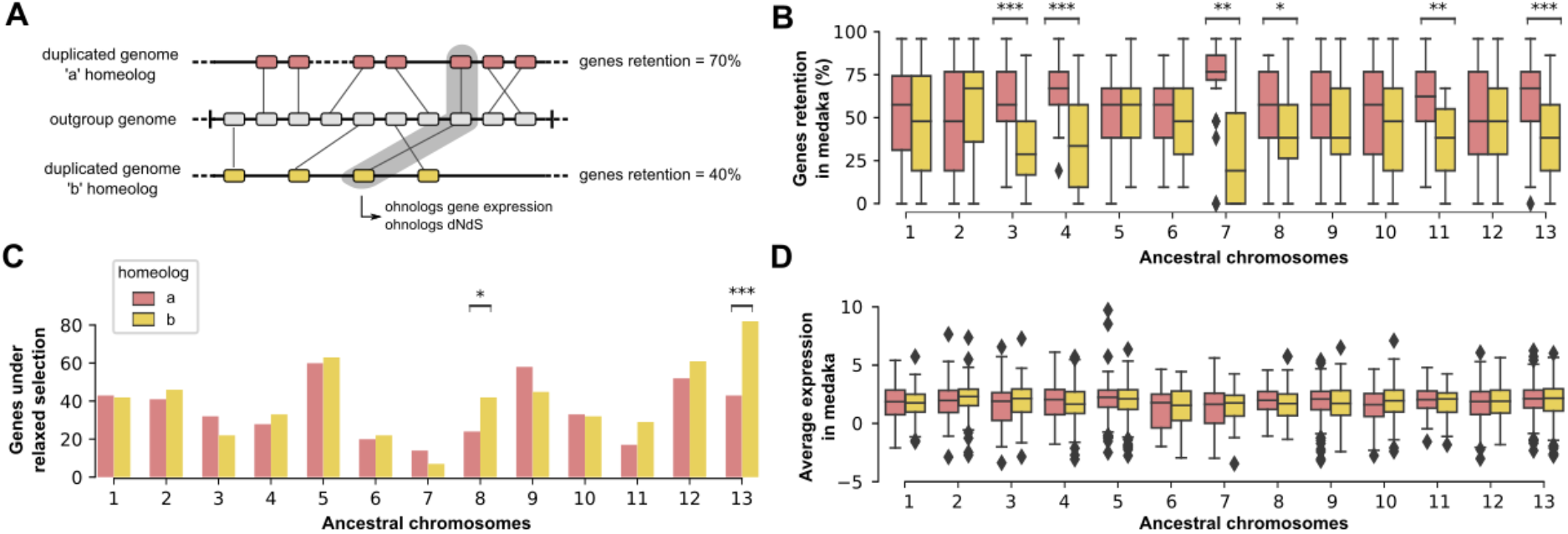
Differences in genes retention, selective pressure and gene expression between duplicated chromosomes. **A**. Schematic example of genes retention calculation. Using an outgroup genome as an approximation of the ancestral gene order, we assess genes retention on each duplicated chromosome in teleost genomes, by 10-gene bins, regardless of their genomic location (Methods). **B**. Gene retention on anciently duplicated chromosome copies in medaka, using the spotted gar genome as a proxy for ancestral gene order. Ancestral chromosomes with a significant bias in gene retention on one of the two copies are highlighted (*** p < 0.001, ** p < 0.01, * p < 0.05, Wilcoxon paired tests with Benjamini-Hochberg correction for multiple testing). **C**. Number of genes experiencing relaxed selection compared to their ohnolog across homeologs (Methods; Fisher’s exact tests with Benjamini-Hochberg correction for multiple testing, p-values as in (B)). **D**. Average expression across tissues in medaka. No significant differences in expression were detected between genes of duplicated chromosome copies (Wilcoxon paired tests with Benjamini-Hochberg correction for multiple testing, at α=0.05).

### Differences in selective pressures and gene expression do not explain the observed bias in genes retention

Previous observations have suggested that biased gene retention in allopolyploid genomes reflects partial epigenetic silencing of one parental subgenome (Freeling et al. 2012; Bird et al. 2021). Genes on silenced chromosome copies are expressed at lower levels, and therefore subjected to lower selective pressures and ultimately to pseudogenization. We therefore investigated whether anciently duplicated chromosomes copies that retained fewer genes have decayed because of unequal selective pressure. We estimated dN/dS ratios for 1,263 pairs of TGD-derived paralogs conserved in at least 40 species, and tested whether ohnologs on one of the two anciently duplicated chromosomes systematically underwent more relaxed sequence evolution (Methods). On ancestral chromosome pairs 8 and 13, genes of one chromosome copy experienced a significant relaxation of selection compared to their ohnologous counterparts (Figure 3C). Genes under relaxed selection were located on the homeolog with lower retention, as predicted if low selective pressure promotes gene loss. However, homeologs with significant differences in selective pressure correspond only to 2 out of 6 chromosomes exhibiting strongly biased genes retention (n = 13; p = 0.1923; Fisher’s exact test), with non-significant differences in selective pressure in the same direction as the gene retention bias for two chromosome pairs (4 and 11) and in the opposite direction for the two others (3 and 7). Thus, differences in selective pressures do not explain the observed bias in gene retention.

In addition, we investigated whether ancestral chromosome copies exhibit differences in gene expression, which would have been established after the TGD and have been maintained since. We find no bias in ohnolog gene expression between ancestral chromosomes from each pair. This result is consistent whether looking at average gene expression across 11 tissues in medaka (Figure 3D), zebrafish (Supplementary Figure S8), or tissue by tissue in either medaka and zebrafish (Supplementary Figure S9-S10).

In conclusion, our findings are consistent with previous reports that gene retention on ancestral chromosomes was biased following the teleost duplication (Conant 2020). However, we find that biased gene retention in teleosts is not correlated with other features classically observed in allopolyploids, and our results suggest that biased gene retention can occur following autopolyploidization, possibly driven by distinct and underappreciated factors.

### The comparative atlas enhances teleost gene and genome annotations

Finally, we investigated how the comparative atlas may improve crucial resources for fish evolutionary, ecological and functional genomics. The Zebrafish Information Network (ZFIN) provides manually curated, high-quality reference annotations for zebrafish genes and implements rigorous conventions for gene naming (Ruzicka et al. 2019). These gene names are then propagated to orthologous genes in other teleost genomes, providing the basis of the entire teleost gene nomenclature and functional annotation transfers. In an effort to convey evolutionary meaning within the gene names, zebrafish paralogs descended from the TGD are identified with an ‘a’ or ‘b’ suffix. ZFIN guidelines recommend that adjacent genes should carry the same suffix when they belong to a continuous syntenic block inherited since the TGD, sometimes called syntelogs (Zhao et al. 2017). This aspiration has however been difficult to implement in the absence of a high-resolution map of zebrafish duplicated regions and their ancestral chromosomes of origin. To assess the consistency in consecutive zebrafish gene names, we extracted and compared zebrafish ‘a’ and ‘b’ gene suffixes along duplicated regions from the comparative atlas (Figure 4). We find that despite previously mentioned efforts, zebrafish ‘a’ and ‘b’ gene suffixes are not consistent with the polyploid history of the zebrafish genome (Figure 4A): 43% of gene suffixes would have to be reassigned in order to reflect the shared history of syntenic genes, which would be impractical to implement (Methods). Gene annotations are therefore unhelpful to study large-scale processes such as chromosome evolution or genome-wide rediploidization. Additionally, the ZFIN nomenclature does not impart a suffix to singleton genes, which correspond to TGD-duplicated genes where one of the copies was eventually lost. As a result, only 26% of zebrafish genes are annotated with an ‘a’ or ‘b’ suffix in ZFIN.

**Figure 4:**
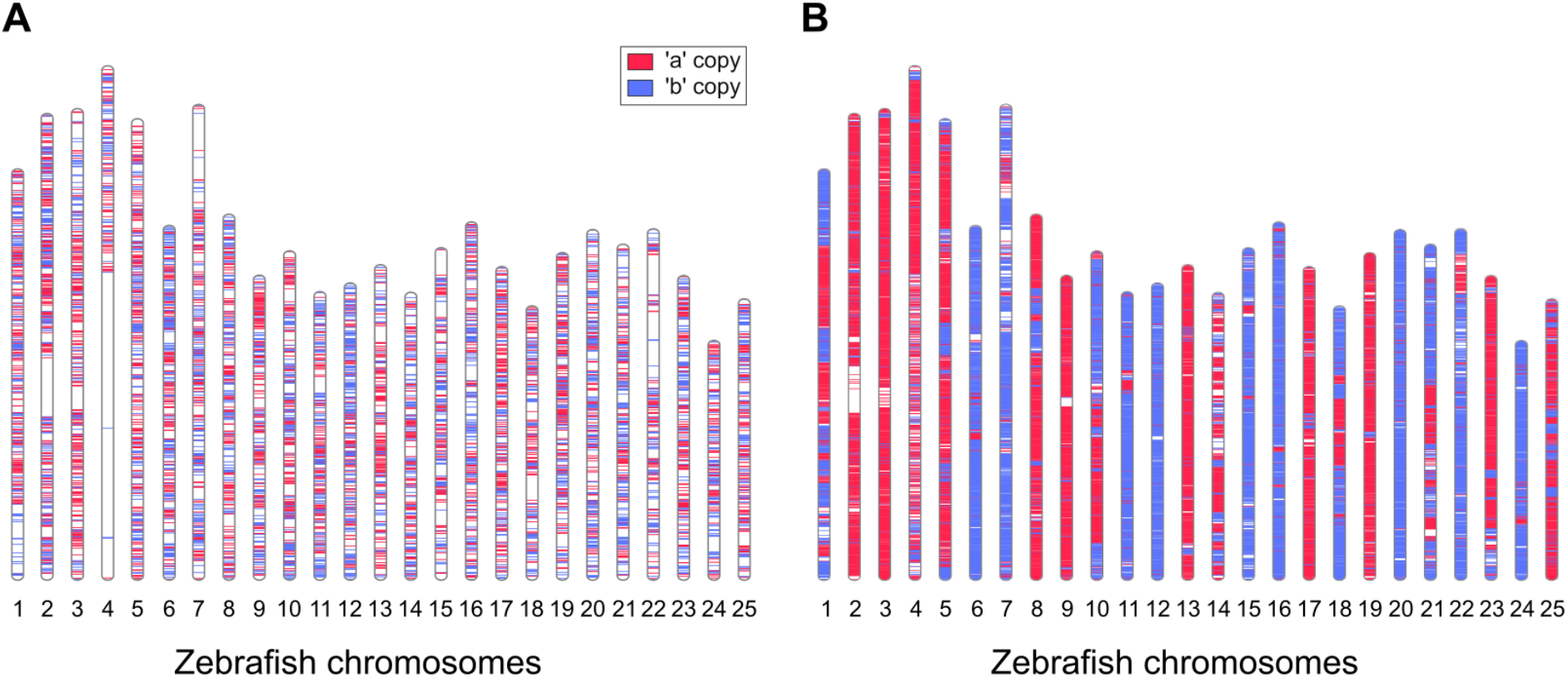
Zebrafish gene names are not evolutionary-consistent. **A**. Karyotypic localization of zebrafish ‘a’ and ‘b’ TGD paralogs, according to the ZFIN annotation. ZFIN does not annotate genes as either ‘a’ or ‘b’ when one of the TGD paralogs has been lost, and these genes are not represented here. **B**. Complementary annotation of zebrafish ‘a’ and ‘b’ gene copies using the comparative atlas (84% of zebrafish genes annotated, including genes without a WGD paralog).

Using the comparative atlas, we annotated which paralogs were retained for 84% of all zebrafish genes, providing a complementary resource that significantly extends the evolutionary annotation of gene histories in this species (Figure 4B). The comparative atlas also includes similar annotations for all 74 studied teleosts, as shown for medaka, stickleback and tetraodon (Supplementary Figure S11). How ancient tetraploids return to a mostly diploid state is an active area of research, where distinguishing which paralog gene has been retained can be essential (Inoue et al. 2015; Robertson et al. 2017; Conant 2020; Simakov et al. 2020; Gundappa et al. 2021). The comparative atlas opens the possibility to formally identify and investigate which ancestral copies have been retained and lost during teleost diversification, and transfer functional gene annotations between model and non-model fish genomes. As such, the comparative atlas constitutes a biologically meaningful, historically accurate insight into reference gene annotations to support further investigations of teleost genome evolution.

## Discussion

Teleost fishes have a long-standing history as tractable model species for vertebrate development and human disease (Ohno et al. 1968; Streisinger et al. 1981; Haffter et al. 1996; Lieschke and Currie 2007; Schartl 2014), and have contributed major breakthroughs in ecological, evolutionary and functional genomics over the years (Braasch et al. 2016; Capel 2017; Xie et al. 2019). Importantly, teleosts have experienced several whole-genome duplications during their diversification, a process which has been essential to early vertebrate evolution. Vertebrates underwent two foundational WGDs over 450 million years ago (Dehal and Boore 2005), and these events have contributed to the establishment of the vertebrate karyotype as we know it (Sacerdot et al. 2018; Simakov et al. 2020), as well as to the expansion of major gene families such as the *Hox* clusters and MHC genes (Castro et al. 2004; Dehal and Boore 2005). As WGDs are rarer in vertebrates than they are in plants, our knowledge regarding post-WGD evolution is anchored on processes observed in plants, where these events are frequent and mechanistically diverse. However, functional constraints on genome and organismal evolution are significantly different between plants and vertebrates. How principles of plant polyploid evolution extend to vertebrates is not well understood, and characterizing ancient WGD events such as the teleost whole-genome duplication is essential to illuminate and interpret the early genetic mechanisms at the origin of vertebrates.

One long-standing question with respect to the teleost duplication is whether the tetraploid teleost ancestor arose via allopolyploidization or autopolyploidization (Martin and Holland 2014; Conant 2020). Previous studies have highlighted the contrasting genomic implications of allopolyploidization and autopolyploidization, where the two subgenomes of allopolyploids are often – although not systematically – subjected to asymmetrical evolution, while in autopolyploids all chromosome copies are highly similar and equally affected by the rediploidization process (Garsmeur et al. 2014; Cheng et al. 2018). We leveraged the comparative atlas to investigate how the post-duplication rediploidization process has shaped extant teleost genomes and reveals the early history of tetraploid fishes. We find evidence for prolonged recombination between entire duplicated chromosomes after the TGD, thus strongly suggesting that the ancestor of teleosts was an autopolyploid. Our results provide the first genome-wide support for delayed rediploidization in teleosts following the TGD, previously suggested as one potential explanation for the enigmatic evolutionary history of the *Hox* and rhodopsin genes in teleosts, and of several gene families in eels (Martin and Holland 2014; Nakamura et al. 2017; Rozenfeld et al. 2019). We demonstrate that delayed rediploidization after the Osteoglossiformes-Clupeocephala lineage split did not only affect these specific gene segments but do extend to entire ancestral chromosomes.

In addition, we find a significant bias in genes retention for a subset of duplicated chromosomes pairs, as also previously observed at a more local scale by (Conant 2020). We lack an explanatory mechanism for these differences, which have been classically linked to allopolyploid genome rediploidization, where one parental subgenome is more expressed, under stronger selective pressure, and retains more genes. Here, we find no clear correlation linking together differences in genes retention, expression level and selective pressure. While the well-studied examples of salmonids and carps genome duplications recapitulate classical models of autopolyploid and allopolyploid rediploidization respectively (Robertson et al. 2017; Xu et al. 2019; Gundappa et al. 2021; Li et al. 2021), polyploid karyotype evolution in vertebrates can also clearly be more complex and involve additional, less appreciated factors that remain to be investigated. Importantly, we highlight that biased gene retention cannot be considered as reliable evidence in favour of allopolyploidization in vertebrates, as we provide formal evidence that the same chromosomes can initially recombine and then experience biased gene retention, suggesting that this bias is unrelated to initial sequence divergence in teleosts.

These novel insights into the rediploidization processes of teleost genomes have important implications for fish evolutionary genomics. First, as the ancestral teleost was a probable autotetraploid, the formal distinction between ancestral subgenomes is irrelevant, and the annotations of chromosomal copies in the comparative atlas should not be mis-interpreted as two distinct parental subsets, where all ‘a’ chromosomes (or ‘b’) descend from a single parental genome. Second, delayed rediploidization has profound consequences on the dynamics of gene evolution. After whole-genome duplication, duplicated gene copies can diverge and undergo specialized evolution by partitioning ancestral functions (sub-functionalization) or acquiring new expression patterns (neo-functionalization) (Ohno et al. 1968; Force et al. 1999; Lynch and Conery 2000). These processes are thought to contribute to diversification and the acquisition of lineage-specific traits: therefore, resolving gene orthology relationships between species is critical to investigate the dynamics of paralog gene retention, divergence and loss and their involvement in phenotypic novelty. However, in this evolutionary scenario, gene paralog sequences only start diverging once recombination is suppressed, which implies that there are no strict 1:1 orthology relationships between duplicated genes across species that separated before meiosis resolution. As such, ‘a’ and ‘b’ gene assignments are not informative of the underlying sequence information contained in genomic regions that maintained meiotic recombination during the early evolution of teleost fishes, and these genes should be considered as tetralogs rather than paralogs and orthologs (Martin and Holland 2014; Robertson et al. 2017). The genetic and functional divergence of these genes has occurred independently in each subsequent lineage, and further inquiry will reveal whether these specific evolutionary dynamics have contributed to lineage-specific evolution in the major teleost clades, as described in salmonids (Robertson et al. 2017; Gundappa et al. 2021). In particular, further work across teleosts and other (paleo)polyploid clades are required to untangle the contributions of mechanistic constraints to meiosis resolution dynamics, and the evolutionary drivers delaying or promoting the genetic divergence of ohnologs.

It is important to note that the teleost comparative atlas comes with a number of limitations that directly stem from the way it was built. First, the assignations to ancestral chromosomes before the TGD are only as good as the state-of-the-art ancestral genome reconstruction. There is a general consensus that the ancestral teleost genome contained 13 chromosomes (Kasahara et al. 2007; Nakatani and McLysaght 2017) – however, the precise delineation of genes descending from each of those 13 groups may be subject to modifications as the field evolves, which may lead to updates in the ancestral assignations of the regions in the comparative atlas. Second, while we show that the comparative atlas is resilient to species-specific errors in ancestral chromosome or gene orthology groups assignations due to the redundant information provided by multiple species, the atlas is ultimately based on gene family tree models, which are sometimes inaccurate or incomplete (Hahn 2007; Som 2015). Inaccurate trees may result in local misspecifications of gene paralogs or ancestral assignations in the comparative atlas. We have previously shown that SCORPiOs significantly improves gene tree accuracy after a WGD event (Parey et al. 2020), and we found only few discrepancies between megabase-scale regions predicted to descend from the same ancestral chromosome from (Nakatani and McLysaght 2017) and paralogy relationships predicted at gene-to-gene resolution by our gene trees. This suggests that the atlas is generally accurate. However, we note that SCORPiOs flagged 2,832 gene trees where sequence identity relationships are inconsistent with their local syntenic context, which may represent areas where the comparative atlas is either less reliable or biologically less informative. To conclude, as teleost fishes have become a high priority taxon for several large-scale projects aiming to extend phylogenetic coverage of vertebrate genome resources (Fan et al. 2020; Rhie et al. 2021), we expect that the genome-scale, clade-wide paralogy and orthology resources we provide here will propel the functional and evolutionary characterization of this major clade encompassing more than half of all vertebrate species.

## Methods

### Libraries and packages

Scripts to build the comparative atlas were written in Python 3.6.8 and assembled together in a pipeline with Snakemake version 5.13.0 (Köster and Rahmann 2012). The ete3 package (version 3.1.1) was used for phylogenetic gene tree manipulation and drawing. Other python package dependencies used for plots and analyses include matplotlib (version 3.1.1), seaborn (version 0.9.0), numpy (version 1.18.4), pandas (version 0.24.2), pingouin (version 0.3.4) for Wilcoxon paired tests and multiple testing correction, and scipy (version 1.4.1) for Fisher’s exact tests. Chromosomes painting and synteny comparisons were drawn with the RIdeogram R package (Hao et al. 2020).

### Genomic resources

Genome assemblies and annotations for the 101 vertebrate genomes were downloaded from various sources, including the NCBI, Ensembl version 95 and GigaDB. The source and assembly version used for each genome are listed in Supplementary Table S1. The gene coordinates files for the 74 teleost genomes in the comparative atlas have been deposited to Zenodo (https://doi.org/10.5281/zenodo.5776772).

### Phylogenetic gene trees

Initial gene trees were built using the Ensembl Compara pipeline (Vilella et al. 2009). Briefly, starting from the sets of proteins derived from the longest transcripts in each genome, we performed an all-against-all BLASTP+ (Altschul et al. 1990), with the following parameters ‘-seg no -max_hsps 1 -use_sw_tback -evalue 1e-5’. We then performed clustering with hcluster_sg to define gene families, using parameters ‘-m 750 -w 0 -s 0.34 -O’. We built multiple alignments using M-Coffee (Wallace et al. 2006), with the command ‘t-coffee -type=PROTEIN -method mafftgins_msa, muscle_msa,kalign_msa,t_coffee_msa -mode=mcoffee’. Next, we conducted phylogenetic tree construction and reconciliation with TreeBeST, using default parameters (Vilella et al. 2009). While TreeBeST remains the most efficient method to build gene trees for large datasets (Noutahi et al. 2016), it systematically infers a number of gene duplication nodes that are overly old and poorly supported (Hahn 2007). We therefore edited the TreeBeST gene trees: we used ProfileNJ (Noutahi et al. 2016) to correct nodes with a very low duplication confidence score (duplication score < 0.1, computed as the fraction of species that retained the genes in two copies after the duplication). Specifically, ProfileNJ rearranges subtrees below these poorly supported node to make them more parsimonious in terms of inferred duplications and losses. Finally, we ran SCORPiOs (version <u>v1.1.0</u>) to account for several whole genome duplication events in the species phylogeny and correct gene trees accordingly: the teleost 3R WGD, using bowfin and gar as outgroups. SCORPiOs reached convergence after 5 iterative rounds of correction. We identified 8,144 teleost gene subtrees out of 17,493 (47%) that were inconsistent with synteny information, of which 5,611 could be corrected (32% of all subtrees). We note that the corrected-to-inconsistent tree ratio (69%) is comparable to our previous application of SCORPiOs to fish data. Similarly, the proportion of errors in initial sequence-based gene trees is in line with our previous application of SCORPiOs to a dataset of 47 teleost species (Parey et al. 2020). We also applied SCORPiOs to correct the nodes corresponding to the carps 4R WGD, using zebrafish as outgroup; and the salmonids 4R WGD, using Northern pike as outgroup. In the presence of LORe in salmonids, as is the case for teleosts, SCORPiOs will attempt to correct gene trees on the basis of synteny information but the correction will be rejected due to being inconsistent with sequence evolution. As a result, the Salmonid 4R WGD will be placed as lineage-specific duplications in LORe trees, consistent with sequence evolution.

### Ancestral gene statistics

We calculated the predicted number of genes in the post-duplication ancestral teleost genome using our set of 26,692 gene trees, and compared this to 60,447 state-of-the art gene trees stored in Ensembl Compara v100. Specifically, to calculate the number of genes inferred in the teleost ancestor (*Osteoglossocephalai*), we counted ancestral gene copies assigned to *Osteoglossocephalai* in the 26,692 and 60,447 TreeBeST phylogeny-reconciled gene trees. This ancestral gene number is an indirect but accurate approximation of the quality of inferred gene trees, since the major challenge is to accurately position duplications at this ancestral node.

### Comparative genomic atlas

The FishComparativeAtlas pipeline annotates teleost genes to post-TGD duplicated chromosomes. It takes as input genomic regions annotated to pre-duplication chromosomes for 4 reference species (zebrafish, medaka, stickleback and tetraodon), and the set of gene trees described above. Segmentation of the four teleost species with respect to the 13 ancestral chromosomes were extracted from (Nakatani and McLysaght 2017) and genomic interval coordinates converted to lists of genes. All genomes were then reduced to ordered lists of genes. We first converted input genomic intervals from zebrafish genome assembly Zv9 to GRCz11 and from medaka genome assembly MEDAKA1 to ASM223467v1, by transferring boundary genes between assemblies using Ensembl gene ID histories. Next, we identified putative WGD sister regions within each of the 4 reference species, as regions sharing a high fraction of TGD-descended paralogs (Supplementary Figure S2B). We grouped regions descended from the same pre-duplication ancestral chromosome, and iteratively annotated pairs of regions into internally consistent sets of ‘a’ and ‘b’ post-duplication sister regions as follows: for each ancestral chromosome, we started from the largest descendant region and arbitrarily defined it as ‘a’. All regions sharing 50% ohnologs or more with this ‘a’ region are identified its ‘b’ paralog(s). Additional search rounds were then conducted to extend the ‘a’ and ‘b’ annotations in a stepwise manner to all regions descended from this ancestral chromosome. The required ohnolog fraction was decreased at each round, and the search was stopped when remaining blocks shared less than 5% ohnologs with previously annotated regions. Since this identification of duplicated regions was performed independently in each of the 4 species, ‘a’ and ‘b’ regions annotations were homogenized to ensure consistency across species (Supplementary Figure S2C). The homogenization step uses orthology relationships from the gene trees and stickleback as an arbitrary guide species: annotations were switched in other species when ‘a’ segments shared more orthologous genes with the ‘b’ region of stickleback. Where conflicts remained for individual genes annotations, we used a majority vote procedure across all four species to define the post-duplication chromosome. Finally, annotations were propagated to all teleost genomes in the gene trees using orthologies.

### Simulation of ancestral chromosome boundary shifts

To simulate incertitude in interval boundaries in the original ancestral genome reconstruction from (Nakatani and McLysaght 2017), we randomly drew new boundaries in the vicinity of their original location according to a Gaussian distribution with standard deviation σ varying in [5, 10, 25, 50, 75, 100] genes. These boundary shifts were independently generated for each of the 4 reference species, with n=100 random noise simulations for each σ value and each reference species. In total, simulations generated 600 noisy input datasets that were processed with the FishComparativeAtlas pipeline in order to assess its robustness to noise. Each of the 600 produced outputs were then compared to the comparative atlas, by counting the proportion of gene families with a reassigned ancestral chromosome (Supplementary Figure S3).

### Early and late rediploidization gene tree topologies

Phylogenetic gene trees were built for the reduced set of 33 genomes as follows. We first filtered the CDS multiple alignments previously computed for the 74 teleosts and 27 non-teleost outgroup genomes set as described above, to retain only genes of all 27 outgroups and 6 teleost genomes, including 3 Osteoglossiformes: paramormyrops (*Paramormyrops kingsleyae*), arapaima (*Arapaima gigas*) and arowana (*Scleropages formosus*) and 3 Clupeocephala: zebrafish *(Danio rerio)*, medaka (*Oryzias latipes)* and stickleback (*Gasterosteus aculeatus)* (Supplementary Figure S5). We used these reduced multiple alignments to build phylogenetic gene trees with TreeBeST (Vilella et al. 2009), using default parameters and the option “-X 10”. This resulted in a set of 14,391 gene trees containing genes of the 33 retained genomes. We then ran SCORPiOs to correct trees for the teleost-specific duplication using the gar and bowfin genomes as outgroups.

To investigate the occurrence of delayed rediploidization, we implemented an extension to the SCORPiOs pipeline (SCORPiOs “LORelEi” for “<u>L</u>ineage-specific <u>O</u>hnolog <u>Re</u>solution <u>E</u>xtension”) to: (i) identify gene trees characterized by sequence-synteny prediction conflicts and (ii) perform likelihood AU-tests (Shimodaira and Hasegawa 2001) to evaluate how AORe and LORe rediploidization tree topologies are supported by gene sequence evolution. For (i), we identify gene trees that SCORPiOs attempts to correct based on syntenic information, but whose correction is rejected due to low sequence-based likelihood. For (ii), we selected the 5,557 gene families containing a gene in at least one reference outgroup (bowfin or gar) and resulting in distinct tree topologies under AORe and LORe. In practice, these topologies can be distinguished when at least one teleost group (Osteoglossiformes or Clupeocephala) retained both ohnologs, although not necessarily in the same species. For each of these 5,557 families, we built 3 gene trees using RaxML 8.2.12 (Stamatakis 2014), with the GTRGAMMA model: the unconstrained maximum likelihood (ML) tree, the constrained AORe topology and the constrained LORe topology (Figure 2B). We then used CONSEL (Shimodaira and Hasegawa 2001) to test for significant differences in log-likelihood reported by RAxML (Stamatakis 2014). A tree topology was rejected when significantly less likely than the ML tree at α=0.05.

We used Circos version 0.69-9 (Krzywinski et al. 2009) to visualize AORe and LORe ohnologs on the medaka genome. Prior to the Circos construction, we used the ‘bundlelinks’ tool available in the circos-tools suite version 0.23 to bundle together consecutive genes with the same rediploidization mode, using 50 genes as the maximum distance parameter (-max_gap 50). Using this setting, isolated AORe and LORe are less visible (i.e have thinner links) than high-confidence regions of consecutive genes displaying the same rediploidization mode. On the Circos, we annotate ancestral chromosomes corresponding to each medaka chromosome with color labels. For clarity purposes, we only add a label if over 17.5% of genes of a given medaka chromosomes are annotated to the ancestral chromosome.

### Functional enrichment tests

For each of the AORe, LORe whole-chromosomes and LORe interspersed sets, we extracted high-confidence ohnologs list in medaka. Specifically, we retained only ohnologs falling in high-confidence AORe and LORe regions defined by “bundled” gene families in the circos representation (i.e., regions formed from neighboring genes with the same rediploidization mode, using 50-genes sized windows). Finally, we used the zebrafish orthologs of these genes (n=248 zebrafish genes for AORe, n=193 for LORe whole-chromosomes and n=215 for LORe interspersed) to perform Gene Ontologies and KEGG pathways enrichment analyses through the WebGestalt web-server (Liao et al. 2019).

### Genes retention on homeologous chromosomes

Because ancestral gene order is particularly difficult to reconstruct in teleosts due to an elevated rate of microsyntenic rearrangements and gene copy losses (Inoue et al. 2015; Nakatani and McLysaght 2017), we take advantage of a non-duplicated outgroup fish genome as a proxy for the ancestral gene order. Here, we make the assumption that consecutive genes on the outgroup genome, all assigned to the same pre-duplication chromosome, represent the ancestral gene order. Using the gene trees, we identify 10,629 1-to-1 orthologies between spotted gar genes and teleost pre-duplication gene families, and 11,599 with bowfin genes. We then used the gene order in the outgroup genomes as an approximation of the ancestral teleost gene order. We reduced outgroup genomes to these genes, and extracted all blocks of consecutive genes annotated to the same pre-duplication chromosome. We computed the percentage of gene copies retained on ‘a’ and ‘b’ homeologs in each extant duplicated genome, using non-overlapping windows of 10 genes along these blocks.

### dN/dS ratio for ohnologous gene copies

We considered 1,263 teleost gene families annotated in the comparative atlas for the dn/ds analysis, selecting all families that contained exactly 2 ohnologous copies in at least 40 teleost genomes (excluding salmonids and carps, which underwent additional WGDs), and exactly one orthologous copy in the bowfin and gar outgroups. We pruned the trees from any species with only one ohnolog copy or where additional gene duplications were present, in order to obtain ‘a’ and ‘b’ clades with the exact same species, for informative dn/ds comparisons. For each gene family, we used translatorX vLocal (Abascal et al. 2010) to (i) translate coding sequences, (ii) align resulting amino acid sequences with Mafft v7.310 (Katoh and Standley 2013) using option ‘--auto’, (iii) trim poorly aligned regions with Gblocks version 0.91b (Castresana 2000) using parameters ‘-b4=2 -b5=h’ and (iv) back-translate the sequences into codon alignments. We used the RELAX model in HyPhy (Wertheim et al. 2015) to estimate dN/dS ratios and test for significant relaxation or intensification of selection on branches of the ‘a’ subtree compared to branches of the ‘b’ subtree. Briefly, RELAX estimates dN/dS distributions across sites on the sets of ‘a’ and ‘b’ branches and fits a selection intensity parameter “k”, which captures the extend of selection intensification (k>1) or relaxation (k<1). Likelihood ratio tests are conducted to compare the alternative model with the k parameter to a null model without. We identified relaxation or intensification of selection on ‘a’ versus ‘b’ branches when the null model was rejected (p-values<0.05, corrected for multiple testing using the Benjamini-Hochberg procedure). Finally, we performed Fisher’s exact tests corrected for multiple testing using the Benjamini Hochberg procedure, to identify chromosome pairs with significantly higher numbers of genes under relaxed evolution on one chromosomal copy.

### Expression level of ohnologous genes

We used RNA-seq datasets across 11 tissues (bones, brain, embryo, gills, heart, intestine, kidney, liver, muscle, ovary, and testis) in zebrafish and medaka from the PhyloFish database (Pasquier et al. 2016), normalized into TPM (transcripts per million transcripts) and quantile normalized across tissues as previously described (Parey et al. 2020). Gene IDs were then converted from Ensembl version 89 to version 95 using conversion tables downloaded from BioMart (Smedley et al. 2009). Average expression across tissue (Figure 3C, Supplementary Figure S8) and by-tissue expression (Supplementary Figure S9-S10) were calculated for ohnologs grouped by their ancestral chromosome of origin.

### ZFIN gene names

Zebrafish ZFIN gene names were extracted using BioMart (Smedley et al. 2009) from the Ensembl database (version 95). We extracted the last letter of gene names, which represents ‘a’ and ‘b’ ohnology annotations in ZFIN. We then computed the minimal number of ‘a’ and ‘b’ ZFIN gene name reassignments that would be necessary to be consistent with the comparative atlas. In the comparative atlas, ‘a’ and ‘b’ labels are arbitrarily assigned to duplicated chromosomes, i.e. genes descended from chromosomes 1a and 1b could be swapped to 1b and 1a. In order to not artificially overestimate discordances, we first swapped such arbitrary ‘a’ and ‘b’ annotations to minimize differences with ZFIN. Finally, we counted the remaining number of ‘a’ and ‘b’ disagreement for zebrafish genes in the comparative atlas that were also annotated in ZFIN.

### Data access

Gene homology relationships and local synteny conservation between teleosts can be interactively browsed through the Genomicus webserver (Nguyen et al. 2022), accessible at https://www.genomicus.bio.ens.psl.eu/genomicus-fish-03.01/cgi-bin/search.pl. The homology relationships from the comparative atlas can be downloaded as flat or HTML files via an ftp server (ftp://ftp.biologie.ens.fr/pub/dyogen/genomicus-fish/03.01/ParalogyMap), along with the gene trees in New Hampshire eXtended (.nhx) format (ftp://ftp.biologie.ens.fr/pub/dyogen/genomicus-fish/03.01/protein_trees.nhx),

## Software availability

All code for the FishComparativeAtlas pipeline is publicly available on GitHub (https://github.com/DyogenIBENS/FishComparativeAtlas). An archive containing a stable version of the code along with all input data (including gene trees) and the final atlas has been deposited to Zenodo (https://doi.org/10.5281/zenodo.5776772), to reproduce the generation of the comparative atlas or directly inspect it. The SCORPiOs LORelEi extension is available on GitHub (https://github.com/DyogenIBENS/SCORPIOS) and has also been deposited to Zenodo (https://doi.org/10.5281/zenodo.6913688), along with all input data, environments and outputs. In addition, both the FishComparativeAtlas pipeline and SCORPiOs LORelEi extension are available in the Supplemental Material.

## Acknowledgements

We thank Pierre Vincens for the coordination of computing resources and all members of the GenoFish consortium for fruitful discussions. We also thank Dan Macqueen and two anonymous reviewers for their critical reading of this manuscript and their helpful suggestions to improve its clarity.

