## Supplementary Material for "An atlas of fish genome evolution reveals delayed rediploidization following the teleost whole-genome duplication"

[3] *Current address*: Institut Pasteur, Université de Paris, CNRS UMR 3525, INSERM UA12, Comparative Functional Genomics group, F-75015 Paris, France.

* Corresponding authors

### Supplementary Material

#### Supplementary Tables

Supplementary Table S1: List of the 101 genome assemblies used in this study.
Supplementary Table S2: Ohnologs retention across the 74 teleost genomes.
Supplementary Table S3: Gene ontology terms enriched in LORe interspersed
Supplementary Table S4: KEGG pathway enriched in LORe interspersed
Supplementary Table S5: Total gene retention on homeologous chromosomes in the post-duplication teleost ancestor *Osteoglossocephalai*.
Supplementary Table S6: Conservative estimate of total gene retention on homeologous chromosomes in Medaka.

#### Supplementary Figures

Supplementary Figure S1: Species tree of the 101 genomes used in this study.
Supplementary Figure S2: Illustration of the key steps in the comparative atlas workflow.
Supplementary Figure S3: Noise robustness of the comparative atlas pipeline.
Supplementary Figure S4: Impact of SCORPiOs on the establishment of the comparative atlas.
Supplementary Figure S5: Karyotypes of medaka, stickleback and tetraodon with 'a' and 'b' gene copies annotations.
Supplementary Figure S6: Species tree of the 6 teleosts for the rediploidisation timing analysis.
Supplementary Figure S7: Gene retention on anciently duplicated chromosome copies in medaka, using the bowfin genome as a proxy for ancestral gene order.
Supplementary S8: Gene retention on anciently duplicated chromosome copies using different ingroup and outgroup genomes.

Supplementary Figure S9: Zebrafish ohnologs gene expression across homeologs, averaged across tissues.
Supplementary Figure S10-S11: Ohnologs gene expression across homeologs in 11 tissues, respectively in Medaka and Zebrafish.

| Species | Source | Data |
|---|---|---|
| *Acanthochromis polyacanthus* | ensembl | ensembl V95 |
| *Amphilophus citrinellus* | ensembl | ensembl V95 |
| *Amphiprion ocellaris* | ensembl | ensembl V95 |
| *Amphiprion percula* | ensembl | ensembl V95 |
| *Anabas testudineus* | ensembl | ensembl V95 |
| *Anolis carolinensis* | ensembl | ensembl V95 |
| *Astatotilapia calliptera* | ensembl | ensembl V95 |
| *Astyanax mexicanus* | ensembl | ensembl V95 |
| *Bos taurus* | ensembl | ensembl V95 |
| *Caenorhabditis elegans* | ensembl | ensembl V95 |
| *Canis lupus familiaris* | ensembl | ensembl V95 |
| *Chrysemys pictabellii* | ensembl | ensembl V95 |
| *Ciona intestinalis* | ensembl | ensembl V95 |
| *Cynoglossus semilaevis* | ensembl | ensembl V95 |
| *Cyprinodon variegatus* | ensembl | ensembl V95 |
| *Danio rerio* | ensembl | ensembl V95 |
| *Drosophila melanogaster* | ensembl | ensembl V95 |
| *Eptatretus burgeri* | ensembl | ensembl V95 |
| *Esox lucius* | ensembl | ensembl V95 |
| *Fundulus heteroclitus* | ensembl | ensembl V95 |
| *Gadus morhua* | ensembl | ensembl V95 |
| *Gallus gallus* | ensembl | ensembl V95 |
| *Gambusia affinis* | ensembl | ensembl V95 |
| *Gasterosteus aculeatus* | ensembl | ensembl V95 |
| *Haplochromis burtoni* | ensembl | ensembl V95 |
| *Hippocampus comes* | ensembl | ensembl V95 |
| *Homo sapiens* | ensembl | ensembl V95 |
| *Ictalurus punctatus* | ensembl | ensembl V95 |
| *Kryptolebias marmoratus* | ensembl | ensembl V95 |
| *Labrus bergylta* | ensembl | ensembl V95 |
| *Latimeria chalumnae* | ensembl | ensembl V95 |
| *Lepisosteus oculatus* | ensembl | ensembl V95 |
| *Loxodonta africana* | ensembl | ensembl V95 |
| *Mastacembelus armatus* | ensembl | ensembl V95 |
| *Maylandia zebra* | ensembl | ensembl V95 |
| *Mola mola* | ensembl | ensembl V95 |
| *Monodelphis domestica* | ensembl | ensembl V95 |
| *Monopterus albus* | ensembl | ensembl V95 |
| *Mus musculus* | ensembl | ensembl V95 |
| *Neolamprologus brichardi* | ensembl | ensembl V95 |

| Species | Source | Data |
|---|---|---|
| *Oreochromis niloticus* | ensembl | ensembl V95 |
| *Oryzias latipes* | ensembl | ensembl V95 |
| *Oryzias melastigma* | ensembl | ensembl V95 |
| *Paramormyrops kingsleyae* | ensembl | ensembl V95 |
| *Petromyzon marinus* | ensembl | ensembl V95 |
| *Poecilia formosa* | ensembl | ensembl V95 |
| *Poecilia latipinna* | ensembl | ensembl V95 |
| *Poecilia mexicana* | ensembl | ensembl V95 |
| *Poecilia reticulata* | ensembl | ensembl V95 |
| *Pundamilia nyererei* | ensembl | ensembl V95 |
| *Pygocentrus nattereri* | ensembl | ensembl V95 |
| *Scleropages formosus* | ensembl | ensembl V95 |
| *Scophthalmus maximus* | ensembl | ensembl V95 |
| *Seriola dumerili* | ensembl | ensembl V95 |
| *Seriola lalandi dorsalis* | ensembl | ensembl V95 |
| *Stegastes partitus* | ensembl | ensembl V95 |
| *Takifugu rubripes* | ensembl | ensembl V95 |
| *Tetraodon nigroviridis* | ensembl | ensembl V95 |
| *Xenopus tropicalis* | ensembl | ensembl V95 |
| *Xiphophorus couchianus* | ensembl | ensembl V95 |
| *Xiphophorus maculatus* | ensembl | ensembl V95 |
| *Acanthopagrus schlegelii* | gigaDB | http://gigadb.org/dataset/100409 |
| *Betta splenden* | gigaDB | http://gigadb.org/dataset/100433 |
| *Channa argus* | gigaDB | http://gigadb.org/dataset/100279 |
| *Eleginops maclovinus* | gigaDB | http://gigadb.org/dataset/102163 |
| *Hippocampus erectus* | gigaDB | http://gigadb.org/dataset/100298 |
| *Lateolabrax maculatus* | gigaDB | http://gigadb.org/dataset/100458 |
| *Protosalanx hyalocranius* | gigaDB | http://gigadb.org/dataset/100262 |
| *Alligator mississippiensis* | NCBI | GCF_000281125.3 |
| *Austrofundulus limnaeus* | NCBI | GCF_001266775.1 |
| *Boleophthalmus pectinirostris* | NCBI | GCF_000788275.1 |
| *Branchiostoma belcheri* | NCBI | GCF_001625305.1 |
| *Branchiostoma floridae* | NCBI | GCF_000003815.1 |
| *Carassius auratus* | NCBI | GCF_003368295.1 |
| *Clupea harengus* | NCBI | GCF_000966335.1 |
| *Columba livia* | NCBI | GCF_000337935.1 |
| *Coturnix japonica* | NCBI | GCF_001577835.1 |
| *Cyprinus carpio* | NCBI | GCF_000951615.1 |
| *Electrophorus electricus* | NCBI | GCF_003665695.1 |
| *Falco peregrinus* | NCBI | GCF_000337955.1 |
| *Larimichthys crocea* | NCBI | GCF_000972845.1 |

| Species | Source | Data |
|---|---|---|
| *Lates calcarifer* | NCBI | GCF_001640805.1 |
| *Nothobranchius furzeri* | NCBI | GCF_001465895.1 |
| *Notothenia coriiceps* | NCBI | GCF_000735185.1 |
| *Oncorhynchus kisutch* | NCBI | GCF_002021735.1 |
| *Oncorhynchus mykiss* | NCBI | GCF_002163495.1 |
| *Oncorhynchus tshawytscha* | NCBI | GCF_002872995.1 |
| *Pangasianodon hypophthalmus* | NCBI | GCF_003671635.1 |
| *Paralichthys olivaceus* | NCBI | GCF_001970005.1 |
| *Rhincodon typus* | NCBI | GCF_001642345.1 |
| *Salmo salar* | NCBI | GCF_000233375.1 |
| *Salvelinus alpinus* | NCBI | GCF_002910315.1 |
| *Sinocyclocheilus anshuiensis* | NCBI | GCF_001515605.1 |
| *Sinocyclocheilus grahami* | NCBI | GCF_001515645.1 |
| *Sinocyclocheilus rhinocerous* | NCBI | GCF_001515625.1 |
| *Xenopus laevis* | NCBI | GCF_001663975.1 |
| *Callorhinchus milii* | NCBI | GCF_000165045.1 |
| *Amia calva* | NCBI | GCA_017591415.1 |
| *Arapaima gigas* | NCBI | GCA_007844225.1 |
| *Oryzias javanicus* | NCBI | GCA_003999625.1 |
| *Perca flavescens* | NCBI | GCA_004354835.1 |

**Supplementary Table S1: List of the 101 genome assemblies used in this study.** The dataset contains genomic resources from Ensembl version 95, NCBI and GigaDB.

| Species | Cohort | Order | Family | Genes in the atlas | Ohnologs | Ohnolog fraction |
|---|---|---|---|---|---|---|
| *Paramormyrops kingsleyae* | Osteoglossomorpha | Osteoglossiformes | Mormyridae | 20,022 | 6,277 | 31% |
| *Scleropages formosus* | Osteoglossomorpha | Osteoglossiformes | Osteoglossidae | 19,757 | 6,464 | 33% |
| *Arapaima gigas* | Osteoglossomorpha | Osteoglossiformes | Osteoglossidae | 17,221 | 5,119 | 30% |
| *Clupea harengus* | Otomorpha | Clupeiformes | Clupeidae | 20,037 | 5,137 | 26% |
| *Sinocyclocheilus anshuiensis* | Otomorpha | Cypriniformes | Cyprinidae | 36,445 | 9,185 | 25% |
| *Sinocyclocheilus grahami* | Otomorpha | Cypriniformes | Cyprinidae | 38,342 | 9,762 | 25% |
| *Sinocyclocheilus rhinocerous* | Otomorpha | Cypriniformes | Cyprinidae | 37,522 | 9,408 | 25% |
| *Cyprinus carpio* | Otomorpha | Cypriniformes | Cyprinidae | 38,952 | 9,326 | 24% |
| *Carassius auratus* | Otomorpha | Cypriniformes | Cyprinidae | 43,853 | 11,014 | 25% |
| *Danio rerio* | Otomorpha | Cypriniformes | Danionidae | 21,496 | 5,362 | 25% |
| *Astyanax mexicanus* | Otomorpha | Characiformes | Characidae | 20,943 | 5,282 | 25% |
| *Pygocentrus nattereri* | Otomorpha | Characiformes | Serrasalmidae | 21,456 | 5,450 | 25% |
| *Ictalurus punctatus* | Otomorpha | Siluriformes | Ictaluridae | 19,725 | 4,414 | 22% |
| *Pangasianodon hypophthalmus* | Otomorpha | Siluriformes | Pangasiidae | 18,783 | 4,179 | 22% |
| *Electrophorus electricus* | Otomorpha | Gymnotiformes | Gymnotidae | 19,059 | 4,639 | 24% |
| *Esox lucius* | Euteleosteomorpha | Esociformes | Esocidae | 20,069 | 5,491 | 27% |
| *Salmo salar* | Euteleosteomorpha | Salmoniformes | Salmonidae | 34,617 | 9,769 | 28% |
| *Salvelinus alpinus* | Euteleosteomorpha | Salmoniformes | Salmonidae | 29,053 | 7,175 | 25% |
| *Oncorhynchus mykiss* | Euteleosteomorpha | Salmoniformes | Salmonidae | 30,196 | 7,989 | 26% |
| *Oncorhynchus kisutch* | Euteleosteomorpha | Salmoniformes | Salmonidae | 29,352 | 7,489 | 26% |
| *Oncorhynchus tshawytscha* | Euteleosteomorpha | Salmoniformes | Salmonidae | 32,997 | 8,316 | 25% |
| *Protosalanx hyalocranius* | Euteleosteomorpha | Osmeriformes | Salangidae | 15,729 | 3,268 | 21% |
| *Gadus morhua* | Euteleosteomorpha | Gadiformes | Gadidae | 17,167 | 3,257 | 19% |

| Species | Cohort | Order | Family | Genes in the atlas | Ohnologs | Ohnolog fraction |
|---|---|---|---|---|---|---|
| *Hippocampus comes* | Euteleosteomorpha | Syngnathiformes | Syngnathidae | 18,089 | 3,705 | 20% |
| *Hippocampus erectus* | Euteleosteomorpha | Syngnathiformes | Syngnathidae | 14,727 | 2,567 | 17% |
| *Boleophthalmus pectinirostris* | Euteleosteomorpha | Gobiiformes | Oxudercidae | 18,598 | 3,993 | 21% |
| *Channa argus* | Euteleosteomorpha | Anabantiformes | Channidae | 16,985 | 3,558 | 21% |
| *Anabas testudineus* | Euteleosteomorpha | Anabantiformes | Anabantidae | 20,561 | 4,787 | 23% |
| *Betta splendens* | Euteleosteomorpha | Anabantiformes | Osphronemidae | 19,023 | 4,375 | 23% |
| *Monopterus albus* | Euteleosteomorpha | Synbranchiformes | Synbranchidae | 19,031 | 4,102 | 22% |
| *Mastacembelus armatus* | Euteleosteomorpha | Synbranchiformes | Mastacembelidae | 19,776 | 4,448 | 22% |
| *Lates calcarifer* | Euteleosteomorpha | Carangaria | Latidae | 21,349 | 4,997 | 23% |
| *Paralichthys olivaceus* | Euteleosteomorpha | Pleuronectiformes | Paralichthyidae | 19,690 | 4,528 | 23% |
| *Scophthalmus maximus* | Euteleosteomorpha | Pleuronectiformes | Scophthalmidae | 18,658 | 4,302 | 23% |
| *Cynoglossus semilaevis* | Euteleosteomorpha | Pleuronectiformes | Cynoglossidae | 18,814 | 4,001 | 21% |
| *Seriola lalandi dorsalis* | Euteleosteomorpha | Carangiformes | Carangidae | 20,996 | 4,969 | 24% |
| *Seriola dumerili* | Euteleosteomorpha | Carangiformes | Carangidae | 20,009 | 4,698 | 23% |
| *Cyprinodon variegatus* | Euteleosteomorpha | Cyprinodontiformes | Cyprinodontidae | 19,635 | 4,231 | 22% |
| *Fundulus heteroclitus* | Euteleosteomorpha | Cyprinodontiformes | Fundulidae | 19,471 | 4,257 | 22% |
| *Xiphophorus maculatus* | Euteleosteomorpha | Cyprinodontiformes | Poeciliidae | 19,912 | 4,421 | 22% |
| *Xiphophorus couchianus* | Euteleosteomorpha | Cyprinodontiformes | Poeciliidae | 17,278 | 3,501 | 20% |
| *Gambusia affinis* | Euteleosteomorpha | Cyprinodontiformes | Poeciliidae | 18,936 | 4,191 | 22% |
| *Poecilia formosa* | Euteleosteomorpha | Cyprinodontiformes | Poeciliidae | 20,316 | 4,519 | 22% |
| *Poecilia atipinna* | Euteleosteomorpha | Cyprinodontiformes | Poeciliidae | 20,189 | 4,397 | 22% |
| *Poecilia mexicana* | Euteleosteomorpha | Cyprinodontiformes | Poeciliidae | 20,470 | 4,477 | 22% |
| *Poecilia reticulata* | Euteleosteomorpha | Cyprinodontiformes | Poeciliidae | 19,441 | 4,273 | 22% |
| *Kryptolebias marmoratus* | Euteleosteomorpha | Cyprinodontiformes | Rivulidae | 19,079 | 4,081 | 21% |

| Species | Cohort | Order | Family | Genes in the atlas | Ohnologs | Ohnolog fraction |
|---|---|---|---|---|---|---|
| *Austrofundulus limnaeus* | Euteleosteomorpha | Cyprinodontiformes | Rivulidae | 17,243 | 3,445 | 20% |
| *Nothobranchius furzeri* | Euteleosteomorpha | Cyprinodontiformes | Nothobranchiidae | 18,308 | 3,800 | 21% |
| *Oryzias javanicus* | Euteleosteomorpha | Beloniformes | Adrianichthyidae | 17,511 | 3,513 | 20% |
| *Oryzias latipes* | Euteleosteomorpha | Beloniformes | Adrianichthyidae | 20,230 | 4,220 | 21% |
| *Oryzias melastigma* | Euteleosteomorpha | Beloniformes | Adrianichthyidae | 19,983 | 4,292 | 21% |
| *Oreochromis niloticus* | Euteleosteomorpha | Cichliformes | Cichlidae | 18,512 | 4,073 | 22% |
| *Haplochromis burtoni* | Euteleosteomorpha | Cichliformes | Cichlidae | 19,880 | 4,519 | 23% |
| *Pundamilia nyererei* | Euteleosteomorpha | Cichliformes | Cichlidae | 19,877 | 4,526 | 23% |
| *Maylandia zebra* | Euteleosteomorpha | Cichliformes | Cichlidae | 22,113 | 4,969 | 22% |
| *Astatotilapia calliptera* | Euteleosteomorpha | Cichliformes | Cichlidae | 21,333 | 4,736 | 22% |
| *Neolamprologus brichardi* | Euteleosteomorpha | Cichliformes | Cichlidae | 19,861 | 4,467 | 22% |
| *Amphilophus citrinellus* | Euteleosteomorpha | Cichliformes | Cichlidae | 19,981 | 4,531 | 23% |
| *Amphiprion ocellaris* | Euteleosteomorpha | Ovalentaria | Pomacentridae | 20,281 | 4,593 | 23% |
| *Amphiprion percula* | Euteleosteomorpha | Ovalentaria | Pomacentridae | 20,274 | 4,687 | 23% |
| *Acanthochromis polyacanthus* | Euteleosteomorpha | Ovalentaria | Pomacentridae | 20,523 | 4,696 | 23% |
| *Stegastes partitus* | Euteleosteomorpha | Ovalentaria | Pomacentridae | 19,908 | 4,651 | 23% |
| *Labrus bergylta* | Euteleosteomorpha | Labriformes | Labridae | 22,332 | 5,065 | 23% |
| *Notothenia coriiceps* | Euteleosteomorpha | Notothenioidei | Nototheniidae | 19,343 | 3,942 | 20% |
| *Eleginops maclovinus* | Euteleosteomorpha | Notothenioidei | Eleginopidae | 16,843 | 3,468 | 21% |
| *Gasterosteus aculeatus* | Euteleosteomorpha | Gasterosteoidei | Gasterosteidae | 17,979 | 3,722 | 21% |
| *Perca flavescens* | Euteleosteomorpha | Percoidei | Percidae | 18,541 | 3,933 | 21% |
| *Lateolabrax maculatus* | Euteleosteomorpha | Acropomatiformes | Lateolabracidae | 16,876 | 3,164 | 19% |
| *Takifugu rubripes* | Euteleosteomorpha | Tetraodontiformes | Tetraodontidae | 18,086 | 3,646 | 20% |
| *Tetraodon nigroviridis* | Euteleosteomorpha | Tetraodontiformes | Tetraodontidae | 17,619 | 3,625 | 21% |

| Species | Cohort | Order | Family | Genes in the atlas | Ohnologs | Ohnolog fraction |
|---|---|---|---|---|---|---|
| *Mola mola* | Euteleosteomorpha | Tetraodontiformes | Molidae | 18,631 | 4,197 | 23% |
| *Acanthopagrus schlegelii* | Euteleosteomorpha | Eupercaria | Sparidae | 16,040 | 3,220 | 20% |
| *Larimichthys crocea* | Euteleosteomorpha | Acanthuriformes | Sciaenidae | 20,033 | 4,712 | 24% |

**Supplementary Table S2: Duplicate gene (ohnolog) retention across the 74 teleost genomes.** Ohnolog fractions are given as the number of ohnologs identified in the comparative atlas divided by the total number of genes annotated in the atlas for that species. Species are ordered based on their phylogenetic position. Cohort, Order and Family are given according to the DeepFin classification.

| GO ID | GO name | Enrich-ment | P-value | FDR | Genes |
|---|---|---|---|---|---|
| GO:0051338 | regulation of transferase activity | 5.724 | 1.00E-06 | 2.84E-04 | ENSDARG00000038524, ENSDARG00000060316, ENSDARG00000038095, ENSDARG00000077226, ENSDARG00000005941, ENSDARG00000038139, ENSDARG00000086778, ENSDARG00000063583, ENSDARG00000077177, ENSDARG00000015803, ENSDARG00000070360, ENSDARG00000070404 |
| GO:0051174 | regulation of phosphorus metabolic process | 3.524 | 6.80E-05 | 0.0097 | ENSDARG00000038524, ENSDARG00000060316, ENSDARG00000038095, ENSDARG00000077226, ENSDARG00000005941, ENSDARG00000038139, ENSDARG00000086778, ENSDARG00000063583, ENSDARG00000077177, ENSDARG00000015803, ENSDARG00000070360, ENSDARG00000070404, ENSDARG00000062277 |
| GO:1902531 | regulation of intracellular signal transduction | 3.286 | 1.38E-04 | 0.0131 | ENSDARG00000060316, ENSDARG00000038095, ENSDARG00000089873, ENSDARG00000077226, ENSDARG00000038139, ENSDARG00000086778, ENSDARG00000063583, ENSDARG00000077177, ENSDARG00000014465, ENSDARG00000015803, ENSDARG00000003313, ENSDARG00000002816, ENSDARG00000075054 |
| GO:0007167 | enzyme linked receptor protein signaling pathway | 3.686 | 3.52E-04 | 0.0250 | ENSDARG00000038524, ENSDARG00000060316, ENSDARG00000038095, ENSDARG00000031751, ENSDARG00000005941, ENSDARG00000104100, ENSDARG00000010785, ENSDARG00000038139, ENSDARG00000086778, ENSDARG00000038067 |

**Supplementary Table S3: Gene ontology terms enriched in LORe interspersed.** Biological process gene ontology enrichment results for the set of n=215 LORe interspersed zebrafish genes.

| KEGG ID | KEGG pathway | Enrichment | p-value | FDR | genes |
|---|---|---|---|---|---|
| dre04350 | TGF-beta signaling pathway | 5.465 | 2.63E-04 | 0.0426 | ENSDARG00000104100, ENSDARG00000102742, ENSDARG00000010785, ENSDARG00000100666, ENSDARG00000061108, ENSDARG00000038067, ENSDARG00000075226 |

**Supplementary Table S4: KEGG pathway enriched in LORe interspersed.** KEGG pathway enrichment results for the set of n=215 LORe interspersed zebrafish genes.

| Ancestral chromosome | 1 | **2** | **3** | **4** | 5 | **6** | **7** | **8** | **9** | 10 | **11** | 12 | **13** |
|---|---|---|---|---|---|---|---|---|---|---|---|---|---|
| Total pre-duplication gene families [1] | 1526 | **1769** | **1440** | **1144** | 1954 | **682** | **664** | **1498** | **1903** | 1320 | **855** | 1975 | **2502** |
| Genes retained on copy 'a' [2] | 961 | **996** | **1057** | **883** | 1326 | **480** | **523** | **1054** | **1264** | 870 | **650** | 1280 | **1815** |
| Genes retained on copy 'b' [3] | 966 | **1234** | **827** | **570** | 1317 | **433** | **371** | **843** | **1143** | 888 | **450** | 1287 | **1450** |
| Fisher's test p-value [4] | 0.88 | **2.87E-16** | **7.25E-19** | **1.91E-41** | 0.88 | **0.0116606074** | **1.41E-18** | **2.74E-15** | **8.83E-05** | 0.63 | **2.06E-23** | 0.88 | **1.51E-26** |

**Supplementary Table S5: Total gene retention on homeologous chromosomes in the post-duplication teleost ancestor *Osteoglossocephalai*.** The table shows the total number of genes retained on each ancestrally duplicated chromosome copy in *Osteoglossocephalai.* Homeologs with a significant bias in genes retention between the two copies are shown in bold (p-value < 0.05, Fisher's test corrected for multiple testing using the Benjamini-Hochberg procedure).

[1] Total *Osteoglossocephalai* gene families for each pre-duplication chromosome

[2], [3] Gene copies retained on chromosomes 'a' and 'b' respectively

[4] Fisher's test p-value for biased gene retention, corrected for multiple testing with the Benjamini-Hochberg procedure.

We note that Fisher's test assumes independence between observations, which may not be verified here if several genes are affected by a single large deletion. The window-based approach in Figure 3B attempts to limit this potential bias.

| Ancestral chromosome | 1 | **2** | **3** | **4** | 5 | 6 | **7** | **8** | 9 | 10 | **11** | 12 | **13** |
|---|---|---|---|---|---|---|---|---|---|---|---|---|---|
| Total pre-duplication gene families (1) | 1277 | **1519** | **1143** | **955** | 1615 | 584 | **618** | **1217** | 1618 | 1098 | **725** | 1661 | **2099** |
| Genes retained on copy 'a' (2) | 718 | **728** | **775** | **706** | 972 | 349 | **440** | **808** | 963 | 640 | **509** | 988 | **1423** |
| Genes retained on copy 'b' (3) | 697 | **974** | **490** | **368** | 920 | 304 | **224** | **525** | 828 | 577 | **294** | 890 | **923** |
| Non-annotated genes (4) | 110 | **173** | **119** | **80** | 169 | 282 | **156** | **112** | 134 | 96 | **67** | 139 | **185** |
| Fisher's test p-value (5) | 0.426 | **5.21E-19** | **9.71E-33** | **1.30E-54** | 0.074 | 0.011 | **1.169E-34** | **2.19E-30** | **3.45E-06** | 0.010 | **9.61E-30** | 0.001 | **5.48E-54** |
| Fisher's test p-value corrected for non-annotated genes (6) | | **0.017** | **8.38E-11** | **1.79E-29** | | | **0.004** | **8.38E-11** | 1.00 | | **3.41E-13** | 0.204 | **3.39E-20** |

**Supplementary Table S6: Conservative estimate of total gene retention on homeologous chromosomes in Medaka.** The table shows the total number of genes retained on each ancestrally duplicated chromosome copy in medaka, using conservative homeologous copy assignments to medaka genes for which the pre-duplication chromosome is known (Nakatani and McLysaght, 2017) but not the post-duplication chromosome. Ancestrally duplicated chromosomes whose retention remained significantly biased towards the same copy after assigning non-annotated medaka genes to the copy with lower retention are shown in bold (p-value < 0.05, Fisher's test corrected for multiple testing using the Benjamini-Hochberg procedure).

(1) Total medaka gene families for each pre-duplication chromosome.

(2), (3) Medaka gene copies retained on chromosomes 'a' and 'b' respectively.

(4) Non-annotated medaka genes: genes where the post-duplication chromosome is unknown.

(5) Fisher's test p-value for biased gene retention, corrected for multiple testing with the Benjamini-Hochberg procedure, but without accounting for non-annotated genes.

(6) Fisher's test p-value for biased gene retention, corrected for multiple testing with the Benjamini-Hochberg procedure and corrected for non-annotated genes. Cells in bold show chromosomes for which the correction did not affect the retention imbalance. Cells are left blank for unbiased chromosomes and for chromosomes where accounting for non-annotated genes changes the direction of the imbalance.

We note that Fisher's test assumes independence between observations, which may not be verified here if several genes are affected by a single large deletion. The window-based approach in Figure 3B attempts to limit this potential bias. We note that Fisher's test assumes independence between observations, which may not be verified here if several genes are affected by a single large deletion. The window-based approach in Figure 3B attempts to limit this potential bias.

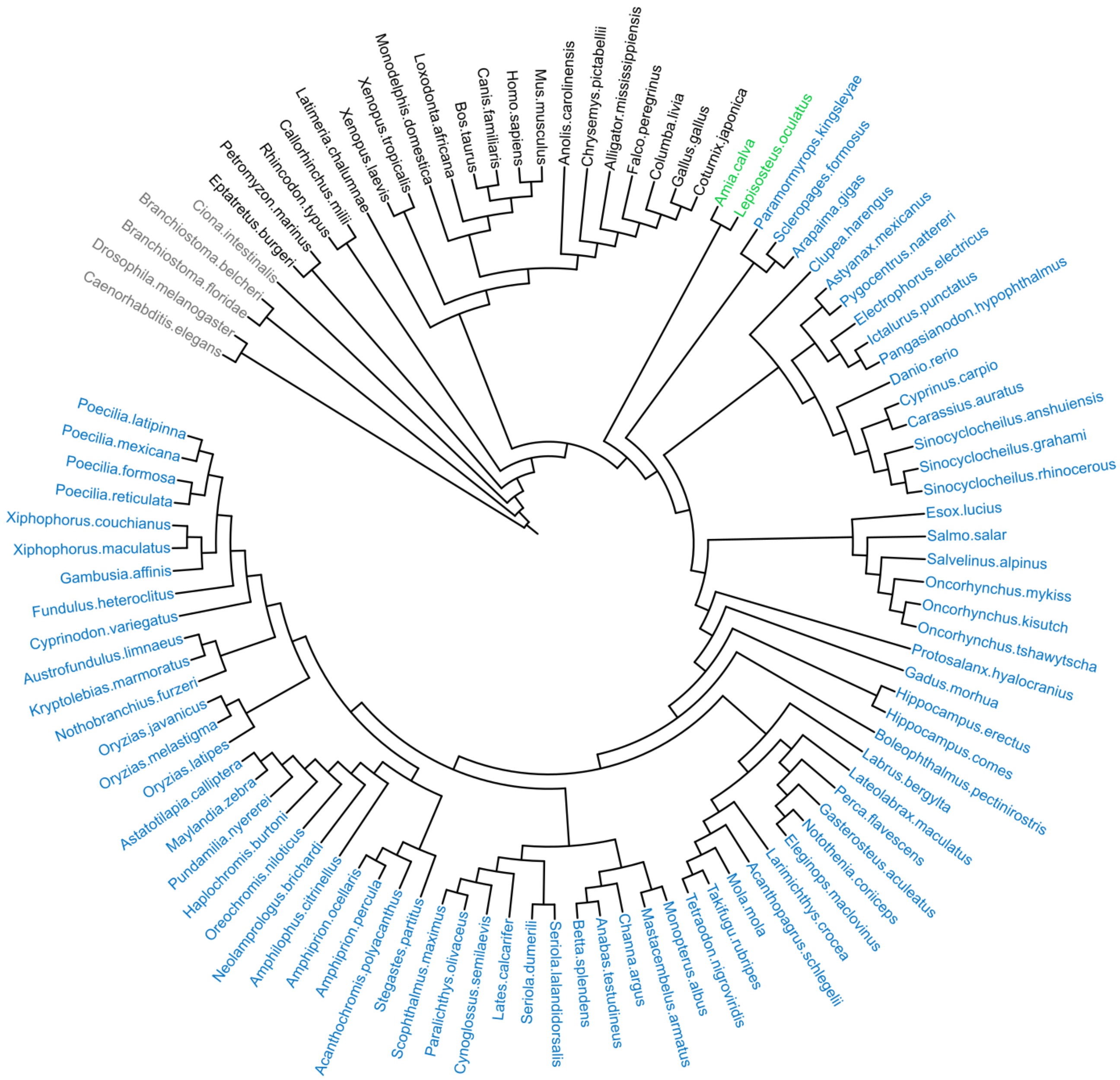


**Supplementary Figure S1: Species tree of the 101 genomes used in this study.** The 74 teleost species are shown in blue, non-duplicated fish outgroups in green, other vertebrates in black and non-vertebrate outgroups in grey. Branch lengths are not to scale.

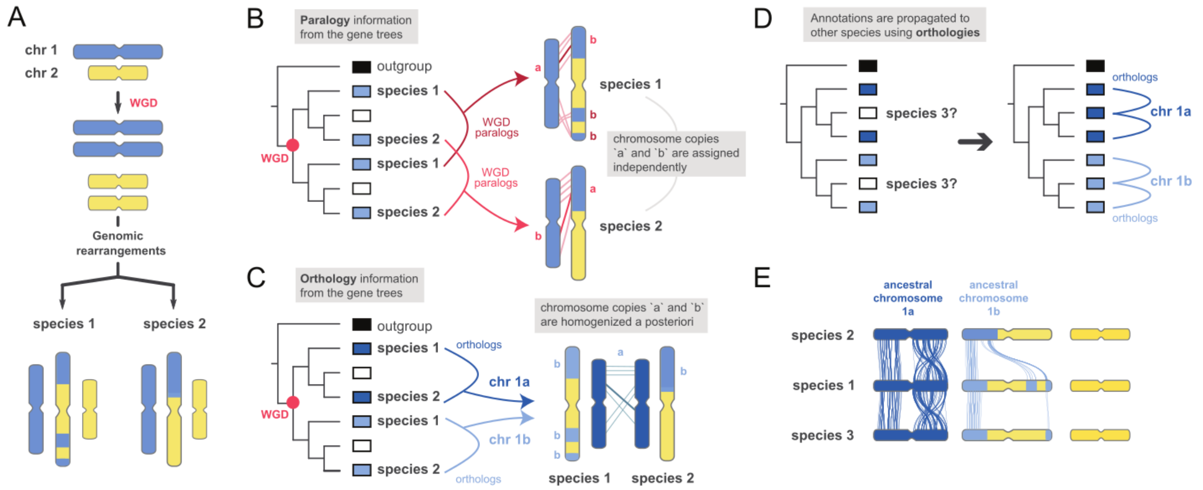


**Supplementary Figure S2: Illustration of the key steps in the comparative atlas workflow. A.** Schematic representation of karyotype histories for two species with a common whole-genome duplication followed by a chromosomal fusion event. **B.** Identification of WGD-duplicated regions using paralogy relationships inferred from gene trees (in red). This identification is performed in each species independently. **C.** Identification of orthologous regions across species inferred from gene trees (in blue). This information is used to homogenize `a` and `b` ancestral chromosome assignations across species. **D.** Propagation of duplicated regions annotations to a non-reference species. Here annotation from reference species 1 and 2 are propagated to species 3 through gene orthologies. **E.** Schematic representation of the comparative atlas. Dark blue and light blue regions correspond to WGD-inherited paralogy regions across all species. Links represent orthologous genes.

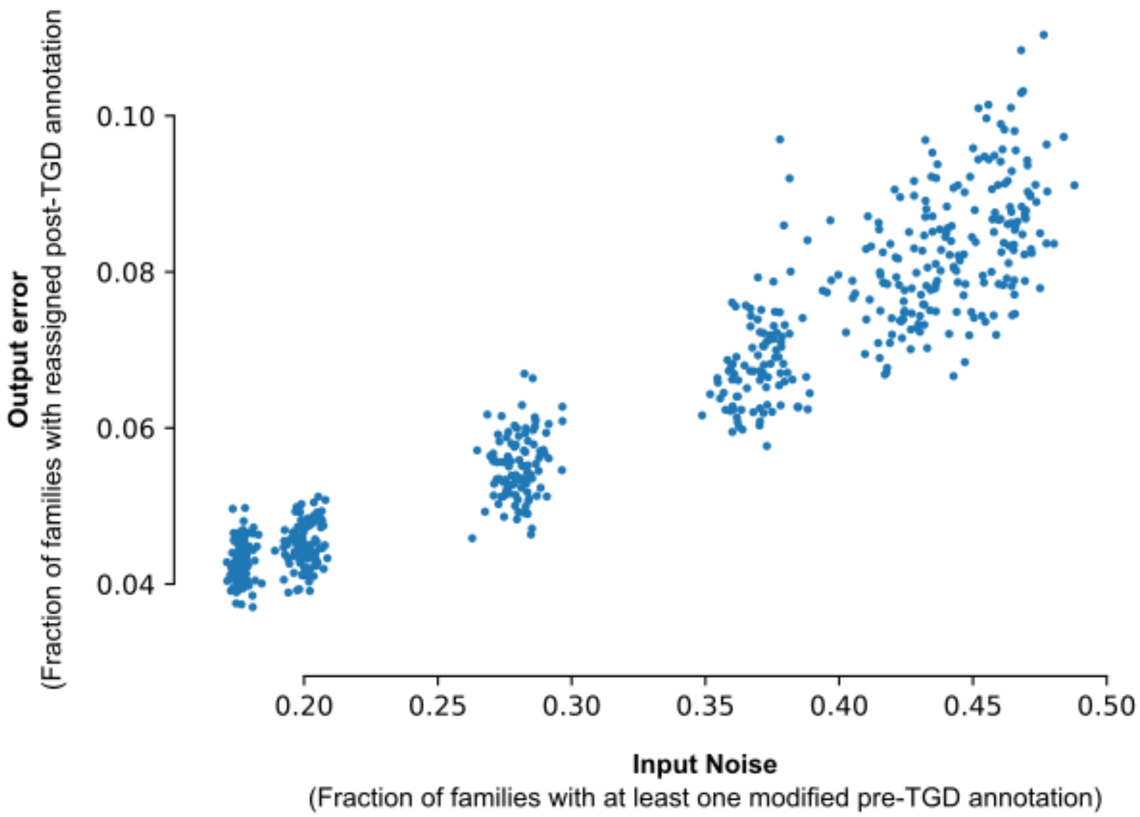


**Supplementary Figure S3: Noise robustness of the comparative atlas.** Proportion of gene families with post-TGD ancestral chromosome reassignments for different input noise settings (Methods). The input noise represents the proportion of families with at least one reference gene having a modified pre-TGD chromosome annotation due to an ancestral chromosome boundary shift in the simulation. The output error represents the fraction of gene families consequently assigned to a different post-TGD ancestral chromosome in the simulation. In the comparative atlas workflow, the majority vote procedure ensures that the majority of these individual errors in inputs are not propagated to the output.

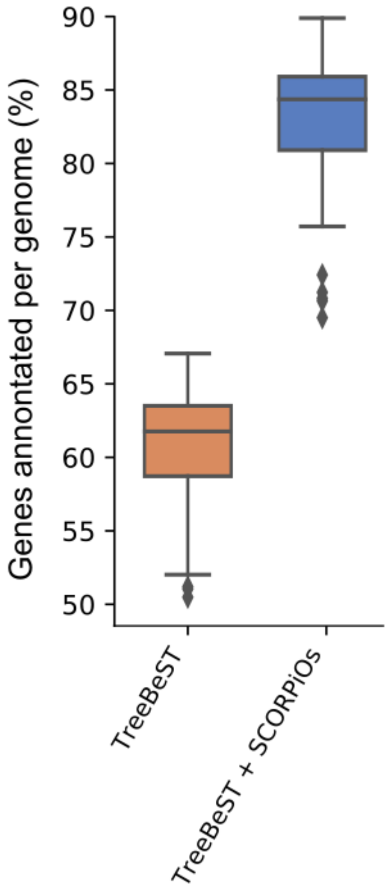


**Supplementary Figure S4: Impact of SCORPiOs on the establishment of the comparative atlas.** Distribution of the proportion of extant genes assigned to an ancestral chromosome in the comparative atlas before (orange) and after (blue) SCORPiOs correction across species. Each boxplot summarizes the distribution of 74 points (one per genome).

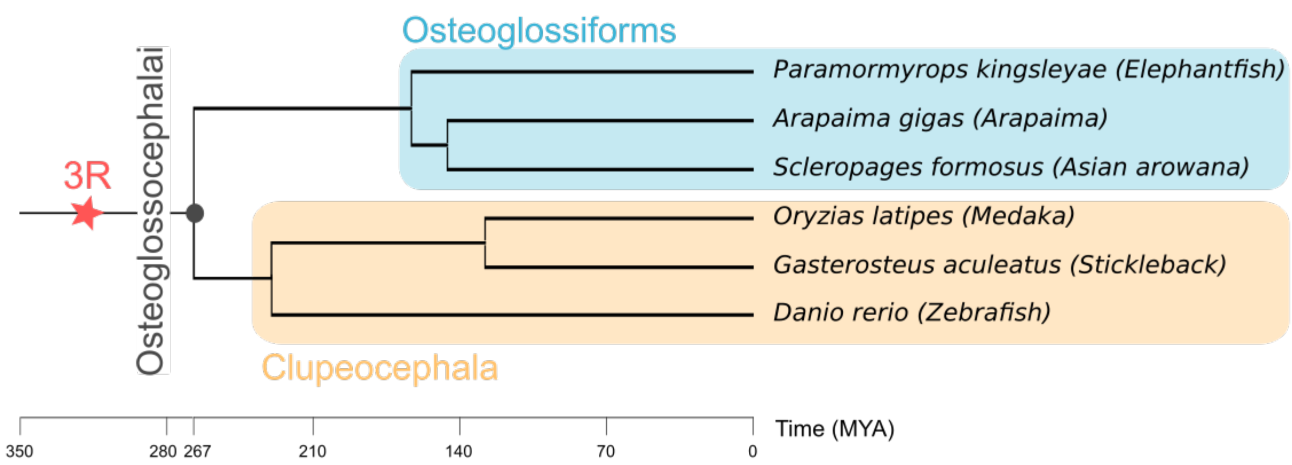


**Supplementary Figure S5: Species tree of the 6 teleosts for the rediploidisation timing analysis.** Divergence times were extracted from TimeTree (Kumar et al. 2017). The teleost duplication is indicated by a red star.

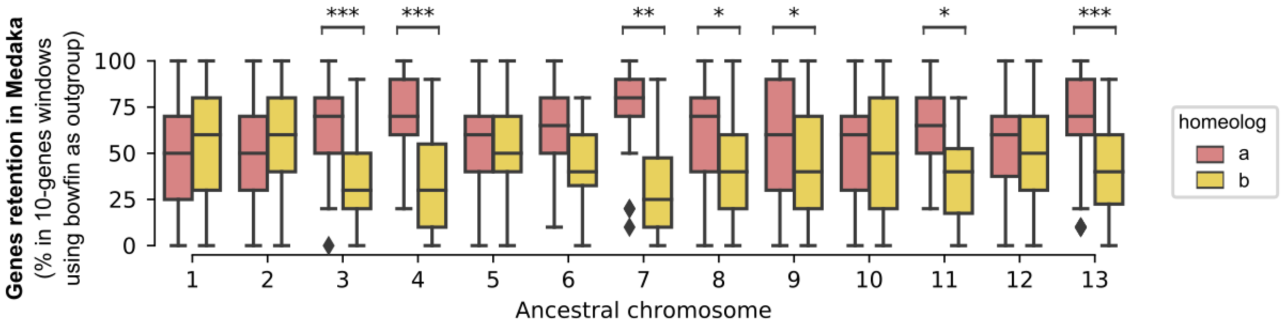


**Supplementary Figure S6: Gene retention on anciently duplicated chromosome copies in medaka, using the bowfin genome as a proxy for ancestral gene order.** Ancestral chromosomes with a significant bias in gene retention on one of the two copies are highlighted (*** p-values < 0.001, ** p-values < 0.01, p-values < 0.05, Wilcoxon paired tests corrected for multiple testing using the Benjamini-Hochberg procedure).

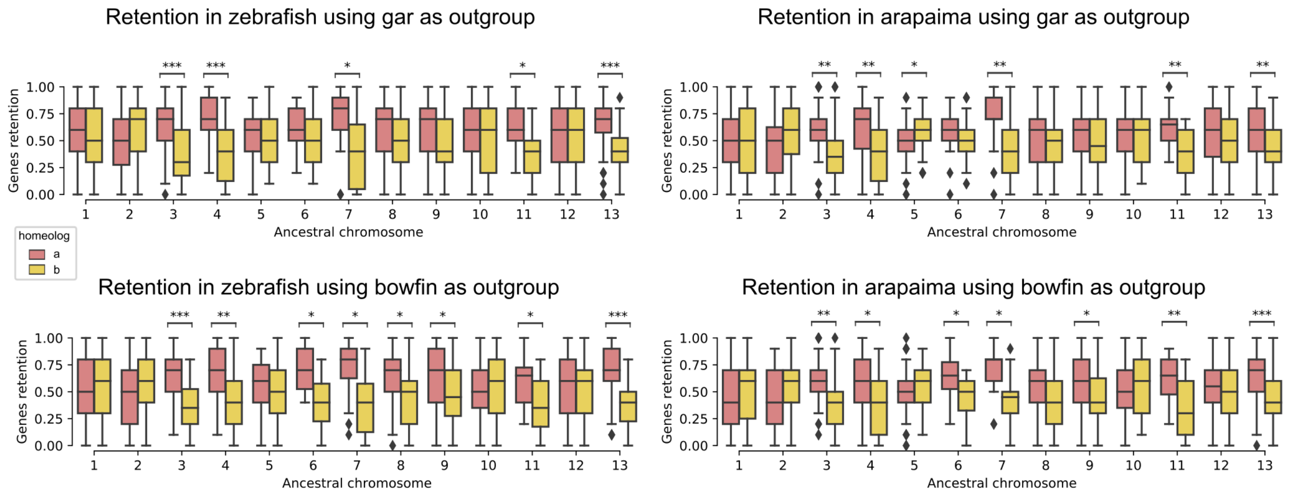


**Supplementary Figure S7: Gene retention on anciently duplicated chromosome copies in zebrafish and arapaima, using the bowfin and the spotted gar genomes as proxies for ancestral gene order.** Ancestral chromosomes with a significant bias in gene retention on one of the two copies are highlighted (*** p-values < 0.001, ** p-values < 0.01, * p-values < 0.05, Wilcoxon paired tests corrected for multiple testing using the Benjamini-Hochberg procedure).

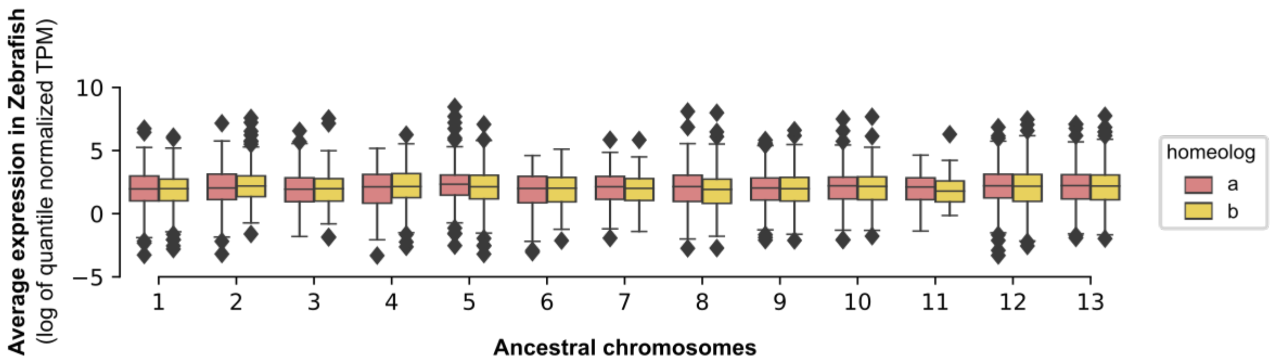


**Supplementary Figure S8: Zebrafish ohnologs gene expression across homeologs, averaged across tissues.** There is no significant difference in expression between genes of anciently duplicated chromosome copies 'a' and 'b' (Wilcoxon paired tests corrected for multiple testing using the Benjamini-Hochberg procedure, at α=0.05).

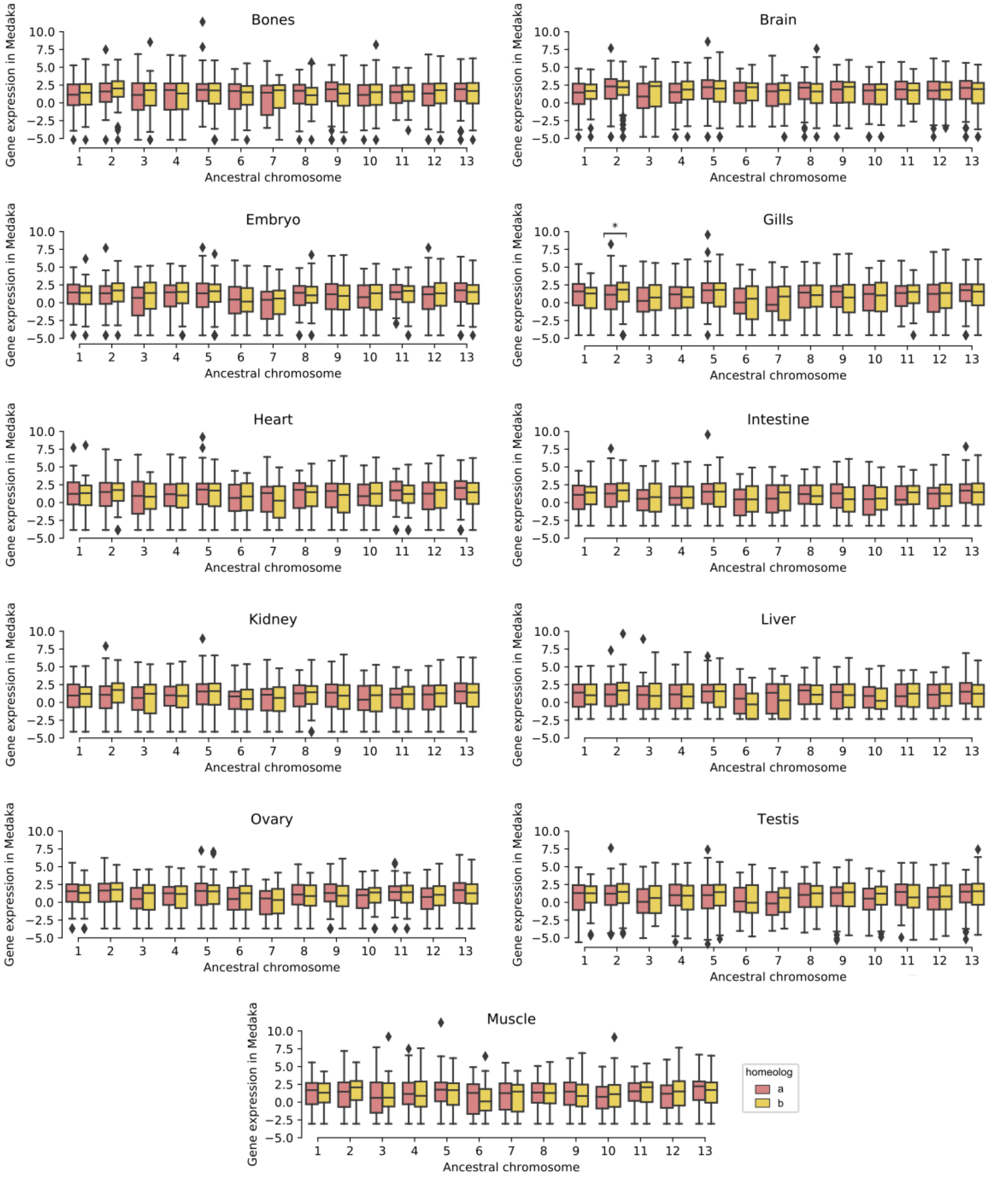


**Supplementary Figure S9: Medaka ohnologs gene expression across homeologs in 11 tissues.** Only for anciently duplicated chromosome 2, in gills, is ohnologous gene expression significantly higher for genes on homeologous copy ‘b’ (* p-value < 0.05, Wilcoxon paired tests corrected for multiple testing using the Benjamini-Hochberg procedure).

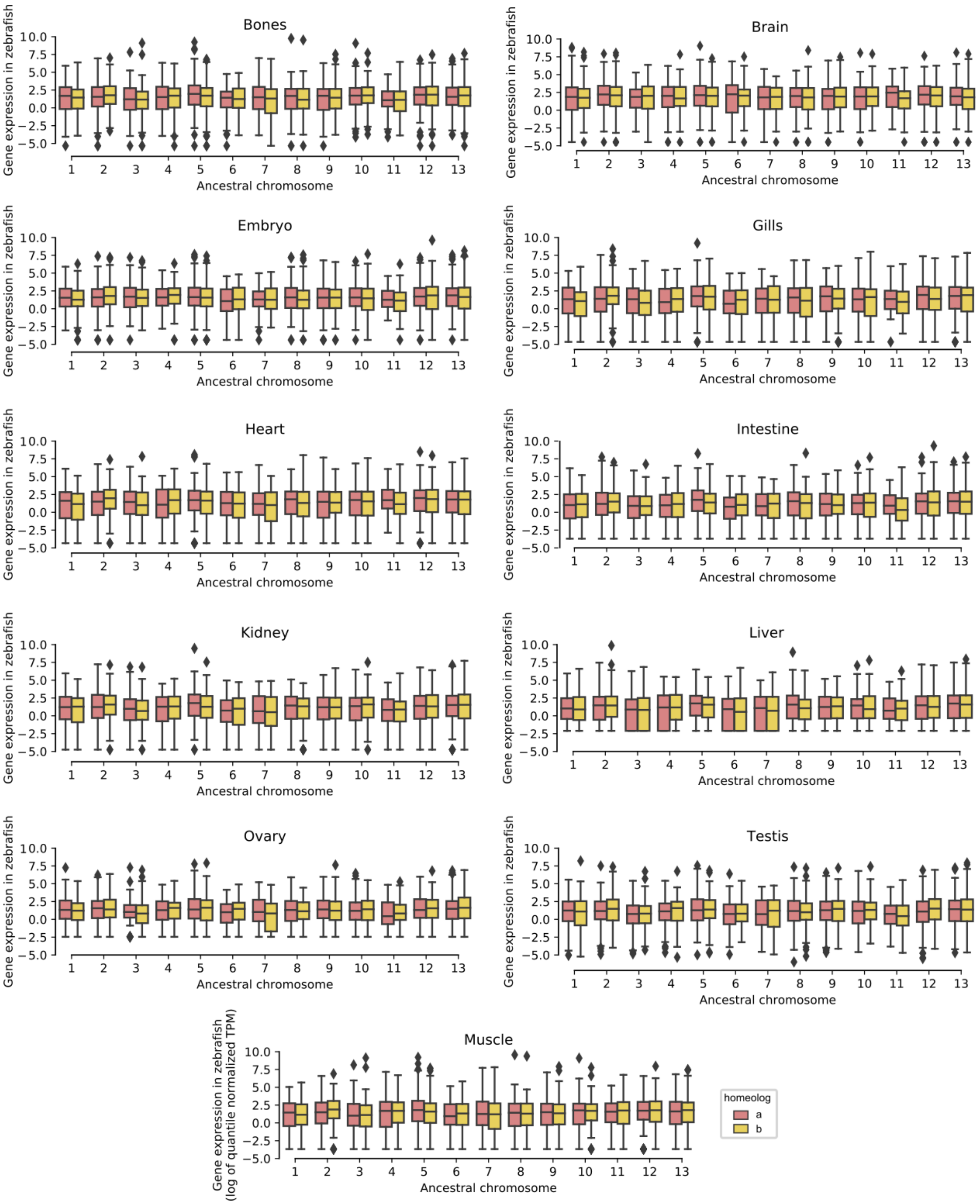


**Supplementary Figure S10: Zebrafish ohnologs gene expression across homeologs in 11 tissues.** There is no significant difference in expression between genes of anciently duplicated chromosome copies 'a' and 'b', in any tissue (Wilcoxon paired tests corrected for multiple testing using the Benjamini-Hochberg procedure, at α=0.05).

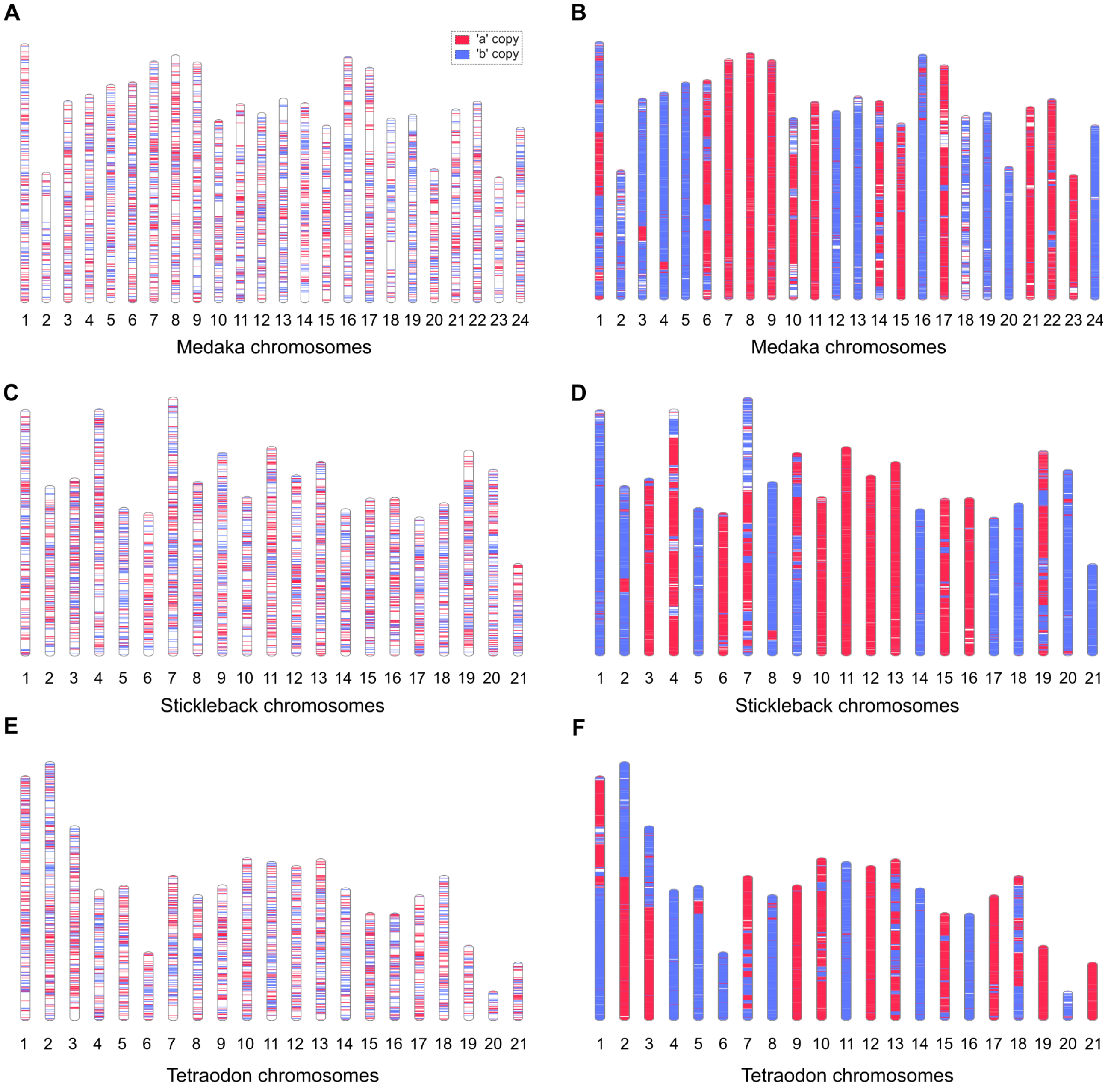


**Supplementary Figure S11: Karyotypes of medaka, stickleback and tetraodon with 'a' and 'b' gene copies annotations. A.** Visualization of medaka 'a' and 'b' copies from ensembl (transfer of zebrafish ZFIN names) on the genome. **B.** Full re-annotation of medaka 'a' and 'b' copies using the comparative atlas. **C.** Stickleback gene copies, as in A. **D.** Re-annotation of stickleback gene copies, as in B. **E.** Tetraodon gene copies, as in A. **F.** Re-annotation of tetraodon gene copies, as in B.
